# A comprehensive survey of developmental programs reveals a dearth of tree-like lineage graphs and ubiquitous regeneration

**DOI:** 10.1101/843888

**Authors:** Somya Mani, Tsvi Tlusty

## Abstract

**Background:** Multicellular organisms are characterized by a wide diversity of forms and complexity despite a restricted set of key molecules and mechanisms at the base of organismal development. Development combines three basic processes — asymmetric cell division, signaling and gene regulation — in a multitude of ways to create this overwhelming diversity of multicellular life-forms. Here, we use a generative model to test the limits to which such processes can be combined to generate multiple differentiation paths during development, and attempt to chart the diversity of multicellular organisms generated.

**Results:** We sample millions of biologically feasible developmental schemes, allowing us to comment on the statistical properties of cell-differentiation trajectories they produce. We characterize model-generated ‘organisms’ using the graph topology of their cell-type lineage maps. Remarkably, tree-type lineage differentiation maps are the rarest in our data. Additionally, a majority of the ‘organisms’ generated by our model appear to be endowed with the ability to regenerate using pluripotent cells.

**Conclusions:** Our results indicate that, in contrast to common views, cell-type lineage graphs are unlikely to be tree-like. Instead, they are more likely to be directed acyclic graphs, with multiple lineages converging on the same terminal cell-type. Furthermore, the high incidence of pluripotent cells in model-generated organisms stands in line with the long-standing hypothesis that whole body regeneration is an epiphenomenon of development. We discuss experimentally testable predictions of our model, and some ways to adapt the generative framework to test additional hypotheses about general features of development.

## Background

Contrary to intuition, the key molecules and mechanisms that go into the development of a human (>200 cell-types [1]) are the same as those required to produce a hydra (just 7 cell-types [2]). More generally, there is a huge diversity of forms and complexity across multicellular organisms, but key molecules of development in *Metazoa* and in multicellular plants are conserved across the respective lineages [3]. The basis of this diversity is illustrated by mathematical models of development which explore possible mechanisms of producing distinctive patterns found in different organisms, for example, segments in *Drosophila* [4], stripes in zebrafish [5], and dorso-ventral patterning in *Xenopus* larvae [6]. At a much broader scale, single cell transcriptomics and lineage tracing techniques have made it possible to map the diversity of forms of extant multicellular organisms [7]. Here, we ask about the *limits of diversity* that development can generate. And reciprocally, we ask what is common among all organisms that undergo development.

Biological development is modular [8], and its outcome rests on gene regulation that is switch-like, rather than continuous [9, 10]. Keeping this in mind, we constructed a generative model of development with three basic ingredients: asymmetric cell division, signaling and gene regulation [11]. Although much is known about the detailed molecular machinery of development [12], naturally, these details come from studies on a few model organisms. We choose to not include all these important particular features in our model for the sake of efficiently and systematically sampling a broad space of developmental schemes. Nonetheless, our model is capable of expressing specific examples of known developmental pathways, which we demonstrate using the *Drosophila* segment polarity network analysed in [9].

We encode organisms in our model as lineage graphs, which show differentiation trajectories of the various cell-types in the organism. Traditionally, mathematical models in the literature elucidate developmental mechanisms responsible for known differentiation trajectories [13]. Here we take the *inverse* approach, and at a much broader scale; we sample across millions of biologically plausible developmental rules and map out the lineage graphs they produce. We purposely do not include selection in the model since it is almost impossible to conjecture and quantify all potential selection factors, such as efficiency, robustness, evolvability, and their intertwined fitness effects on the developmental program. Instead, we sample developmental rules *uniformly* to provide an extensive chart of all possible programs without weighing their relative advantage. Our approach allows us to identify emergent properties that arise from combinations of the ingredients of biological development. We anticipate that such properties are likely to be universal, regardless of the selective pressures faced by these organisms.

By tuning just three biologically meaningful parameters — which control signaling, cellular connectivity and cell division asymmetry — our model produces a rich collection of organisms with diverse cell-type lineage graphs, ranging from those with a single cell-type, to organisms with close to a hundred cell-types. Given the coarse-grained nature of the model, we do not expect model-generated organisms to resemble real organisms in all aspects. Instead, we examine and find hallmarks of multicellular organisms which originate from the fundamental features of development included in the model.

Notably, tree-like lineage graphs are rare in our model. This could indicate that, contrary to popular belief, lineage graphs of real organisms are not tree-like; they are more likely to be directed acyclic graphs (DAGs). Additionally, an unanticipated outcome of our model is that most organisms we generate are capable of whole body regeneration. Our result supports the hypothesis that regeneration is an epiphenomenon of development, rather than a function that evolved separately [14]. The model also produces concrete predictions, and we discuss how these predictions can be experimentally tested on animals like *Planaria* and *Ascidia*, which are well-known models of animal regeneration [15, 16].

### Generative model of development

Organisms in the model contain genomes with *N* distinct genes. By ‘Genes’, we refer not to single genes, but to gene regulatory modules that control cellular differentiation [17]. These genes encode for cell-fate *determinants*. In different cell-types of an organism, different sets of determinants can be present (1) or absent (0). We represent a cell-state as a *N*-length binary string. For example, for *N* = 3, a cell in state *C* = [101] contains determinants 1 and 3 but not the determinant 2. (In Additional file 1: section 1.1 we demonstrate how we can also use ‘determinants’ to encode spatial information using the well-known *Drosophila* segment polarity network as an example.)

Cell-types are ordered according to standard binary ordering, i.e., the cell [101] can equivalently be written as *C*_5_. We only look at whether a given cell-type is present or absent in organisms, rather than the number of cells of any given cell-type. Therefore, since each of the *N* determinants can be either 1 or 0, there are at most 2^*N*^ distinct cell-types in an organism, and 2^2^*N*^^ cell-type compositions for organisms (Fig.1(A)). Note that the number of distinct organisms is larger than 2^2^*N*^^, since different organisms may have the same set of cell-types but distinct lineage graphs (Fig.1(G)).)

**Figure 1.**
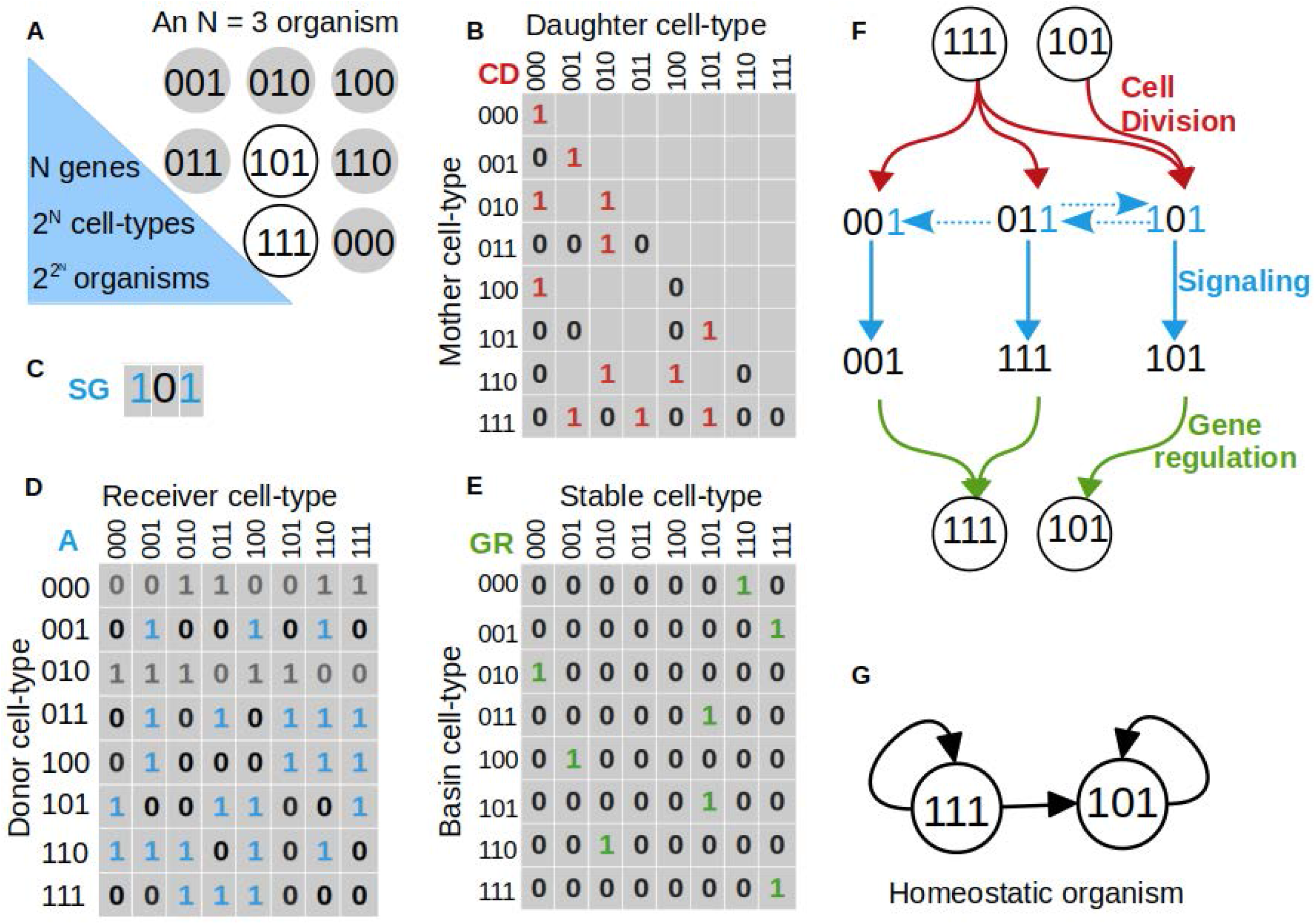
Generative Model. **(A)** An organism with N=3 genes and two cell-types. Circles represent all possible cell-types. The organism is composed of cell-types represented by white circles, and does not contain the grey cell-types. Binary strings written inside the circles represent the presence (1) or absence (0) of determinants in those cell-types. **B,C,D** and **E** describe the rules for development of the organism in **A**. **(B)** Cell division matrix CD. In the model, daughter cell-types cannot contain determinants not present in the mother cell-type, therefore in the figure, such positions in CD are represented by empty boxes. For all *j* such that CD(*i, j*) = 1, cell-type *i* produces cell-type *j* upon cell-division. **(C)** Signaling matrix SG. determinants 1 and 3, which are labelled in blue, act as signaling molecules. **(D)** Signaling adjacency matrix *A. A*(*i, j*) = 1 implies **I** that cell-type *j* receives all signals produced by cell-type i. The rows for cell-types [000] and [010] are greyed out, since these cells produce no signaling molecules **(E)** Gene regulation matrix GR. GR(*i, j*) = 1 implies that cell-type *j* is a stable cell-type, and cell-type *i* maps to cell-type *j*. **(F)** Schematic of ‘organismal development’ in the model. All cell-types synchronously undergo cell-division according to CD, the daughter cells exchange signals according to SG and A, and cells respond to signals through gene regulation according to GR. The process repeats until it reaches a steady state. Here we show how the homeostatic organism in **A** is obtained using the developmental rules matrices in **B,C,D** and **E**. **(G)** Lineage graph of the homeostatic organism in **A**.

We represent development as a repeated sequence of cell division, intercellular signaling, and gene regulation:

#### Cell division

Cells in the model undergo asymmetric cell-division, where daughter cells inherit determinants from the mother cell in an asymmetric manner. That is, a determinant that is present in the mother cell may not be inherited by all its daughter cells due to unequal or insufficient partitioning during division [18]. In the model, asymmetry of cell division is controlled by the parameter *P*_asym_ ∈ [0,1], which is the probability that a daughter cell does not inherit a given determinant from the mother cell; *P*_asym_ = 0 implies symmetric division, where all daughter cells inherit all determinants from the mother cell, and at *P*_asym_ = 1, no daughter cell inherits any determinants from the mother cell.

Although in real multicellular organisms, a single cell only divides into two daughter cells, a single cell-type may represent a population of cells, which need not all behave in the same way [19, 20]. We capture this heterogeneity by allowing cells in our model to divide into more than two types of daughter cells. For any given organism in the model, we predetermine the sets of daughter cells produced by any cell-type randomly according to *P*_asym_, and encode this in a binary matrix CD (Fig.1(B)).

In real organisms, asymmetrically segregating determinants actively influence functionality of cells. Some determinants modulate the response of cells to signals, most famously, the protein Numb, which is an inactivator of Notch signaling, is asymmetrically segregated during the division of neural, muscle and hematopoeitic stem cells [18]. Other asymmetrically segregating proteins act as signaling molecules themselves, for example, the protein Dll1 segregates asymmetrically during neural stem cell division, and is sent as a signal to neighbouring quiescent neural stem cells [21].

#### Signaling

The number of distinct signaling molecules in an organism is controlled by the parameter *P*_sig_ ∈ [0,1], which is the probability that any determinant is a signaling molecule. Parameter *P*_adj_ ∈ [0,1] controls signal reception; a cell-type *C_i_* can receive signals from a cell-type *C_j_* with probability *P*_adj_. Additionally, during simulations, cells can only receive signals from other cell-types present in the same time step, and recipient cells receive all the signals produced by donor cells. As in the case of cell-division, for each organism, the set of signaling molecules, and the pairs of cells that are allowed to exchange signals are predetermined and stored in a binary vector SG, and a binary matrix *A*, respectively (Fig.1(C,D)).

#### Gene regulation

The combination of determinants inherited by a daughter cell during cell-division, and those received as signals from other cells present in the same time-step, together regulate gene expression in this cell. We model gene regulation as random Boolean networks (RBNs) [22]; here, the transcriptional states of genes depend on each other through arbitrarily complex Boolean rules. Updates in gene states lead to updates in the states of determinants, which in turn result in updates in cell-types. In this scheme, some cell-states update to themselves (stable cell-type), and other cell-states ultimately update to one of the stable cell-states, that is, they lie in the basin of a stable cell-type. Note that a cell-type in the model need not be a fixed point (single cell-state) of the gene regulatory network, it can also be an oscillation (multiple cell-states) [23]. In the latter case, the cell-type is represented by all cell-states that are part of the oscillation. Here, we are only concerned with the set of cell-states in the stable state, and not with the sequence of cell-states in oscillations.

In the model, instead of encoding RBNs explicitly, we describe gene regulation directly as the set of stable cell-types and their basins. Stable cell types and cell-states in their basins are both chosen uniform randomly, as described in detail in Methods. Generally, gene regulatory networks with higher *N* have more stable cell-types, but basin sizes, on average, remain small (1-2 cell-states) across all *N* (Additional file 1:Fig.S3). For each organism, we predetermine its gene regulation and encode it in a binary matrix GR (Fig.1(E)). The matrices CD, SG, *A* and GR should be viewed as summaries of all processes that determine cell-fate during the life-time of an organism, and a simulation of the model represents a recapitulation of all these events. Our model is deterministic; for a given organism, the matrices CD, SG, *A* and GR are independently generated, and they remain fixed for the rest of the simulation. The model is also synchronous; all cell-types in the organism divide simultaneously, after which the developing organism is composed only of all daughter cells produced in this step. These daughter cells simultaneously exchange signals, in response to which the states of all the determinants, in each daughter cell are updated simultaneously according to GR (Fig.1(F)). A time-step in the model represents a single repeat of cell-division, signaling and gene regulation.

The process of development ends when the set of cell-types in a developing organism repeats itself. We call this set of cell-types the steady state of the organism, and the number of time-steps between two repeats the period of the steady state. Since this is a finite and deterministic system, starting from any initial condition, such a steady state can always be reached. We call period-1 steady states *homeostatic organisms* (Fig.1(F)). Although organisms with complex, period >1 life-cycles, such as land plants with alternation of generation [24] exist in nature, in this study, we focus on homeostatic organisms. We represent homeostatic organisms as their *cell-type lineage graphs* (Fig.1(G)). The nodes of this graph represent cell-types in the homeostatic organism, and directed edges represent lineage relationships between these cell-types. Let some cell-types *A* and *B* in a homeostatic organism be represented by nodes *V_a_* and *V_b_*, respectively, in its lineage graph. Then, there is an edge from *V_a_* to *V_b_* if one of the daughter cells of *A* gives rise to *B* after one round of cell-signaling and gene regulation.

In the model, all the cell-state transitions that lead to the homeostatic organism starting from an initial cell-type represent the process of development, while the homeostatic organism itself represents the product of development. In this work, we study properties of these homeostatic organisms and their lineage graphs.

### Results

#### Homeostatic graphs span a large range of sizes

We looked at millions of homeostatic organisms, spanning systems with *N* = [3, 4, 5, 6, 7] number of genes (Additional file 1:Fig.S3(A)). 99.88% of these homeostatic organisms had lineage graphs with a single connected component. In the following, we describe lineage graphs of these single-component homeostatic organisms. While a majority of graphs in our data are small (1-5 cell-types), the largest graphs have 89 cell-types (Fig.2(B), Additional file 1:Fig.S4(A)). Across different organisms, number of cell-types in lineage graphs increases with the diversity of daughter cell-types produced (Figs.Additional file 1:Fig.S4(B,C)), and decreases with the level of signaling (Additional file 1:Fig.S4(D,E,F)). Naturally, the number of edges in lineage graphs increases with the number of cell-types, but this increase is slower than that expected for simple random graphs (Additional file 1:Fig.S5).

**Figure 2.**
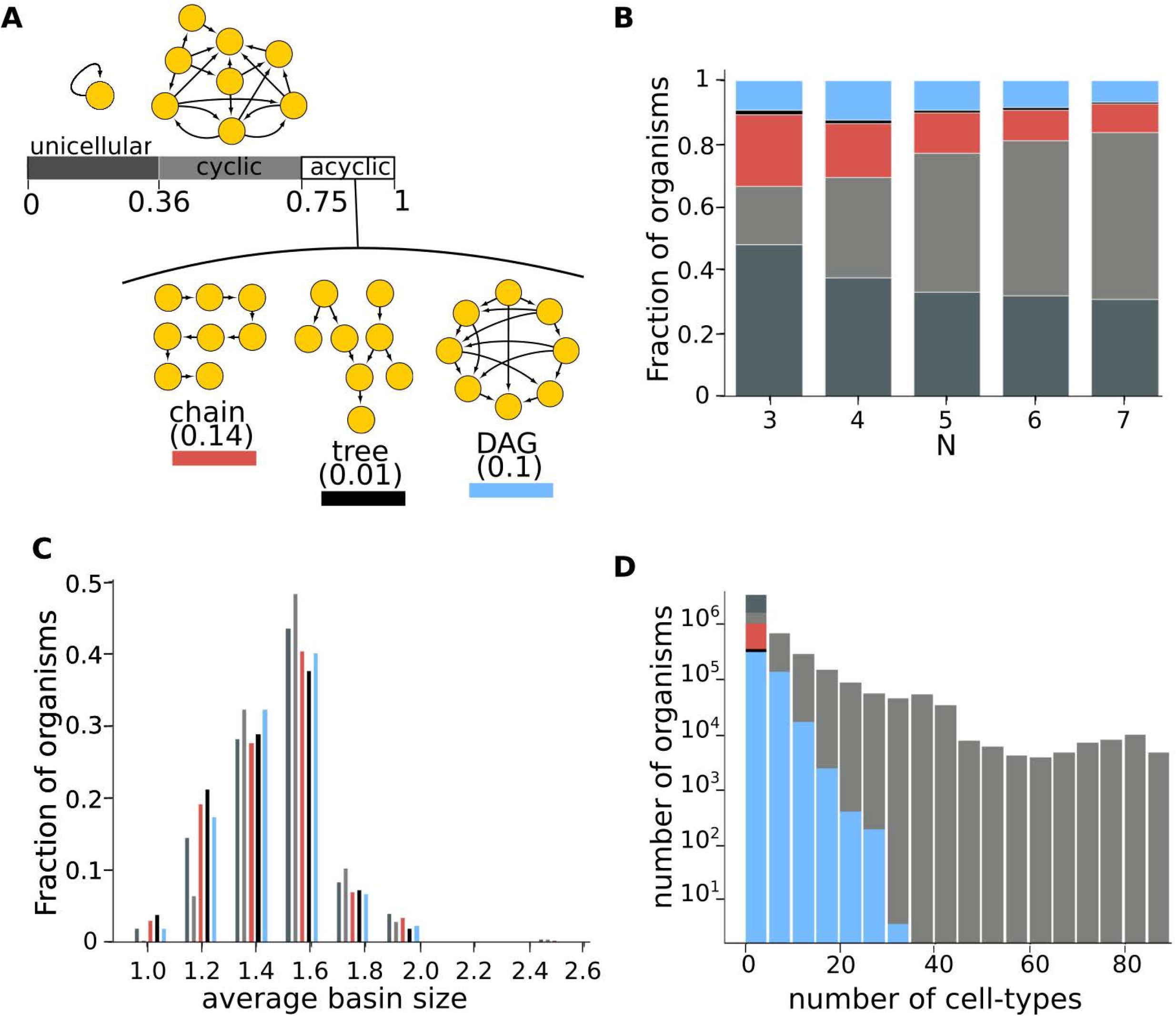
Diversity of lineage graphs. 4852994 graphs were used to generate the following figures. **(A)** Abundance of lineage graph topologies. The bar represents the fraction of graphs that have a given topology. Graphs given along the bar are examples of model-generated lineage graphs. Parameters (*N, P*_asym_, *P*_sig_, *P*_adj_) used to generate these graphs: unicellular:(3, 0, 0, 0), cyclic:(4, 0.2, 0, 0.5), chain:(5, 0, 0.4, 0.9), tree:(6, 0, 0.4, 1), DAG:(4, 0.1, 0, 0.4). Colours indicate lineage graph topology: unicellular:dark grey, cyclic:light grey, chain:red, tree:black, DAG:blue. **(B)** Stacked histogram for topologies of lineage graphs obtained with different N. Heights of colored blocks represent the proportions of corresponding topologies. **(C)** Distribution of basin sizes in the gene regulatory networks across different lineage graph topologies. For any given topology, height of bars indicates the fraction of lineage graphs of that topology that were obtained using a gene regulatory network whose average basin size is given along the horizontal axis. **(D)** Stacked histogram showing distribution of number of cell-types in homeostatic organisms with lineage graphs of various topologies. See also Additional file 1: Fig.S9.

#### Diversity of lineage graph topologies and the dearth of tree-like lineage graphs

Paths in a lineage graph represent differentiation trajectories of the organism’s cell-types. Here, we classify lineage graphs into five topologies, each of which contain qualitatively different paths: (i) unicellular graphs, (ii) cyclic: multicellular graphs that contain some cyclic paths, (iii) chains: acyclic graphs with no branches, (iv) trees: acyclic graphs with branches, and (v) other directed acyclic graphs (referred to here simply as DAGs): acyclic graphs with branches and *links*, which are edges that connect different branches. Links represent the convergence of multiple cell-lineages to the same terminal cell-type. We ignore self-edges during lineage graph classification.

In our data, unicellular graphs are the most abundant (36%). Acyclic graphs (chains, trees and DAGs) comprise about 25% of our graphs. Of these, trees are the rarest (1% across all graphs) and chains are the most abundant (14.3% across all graphs) (Figs.2(A,B)). We find that even after including ‘acyclized’ versions of cyclic graphs in our analyses – i.e. where we merge all nodes belonging to each strongly connected component into single nodes –, trees are still the rarest graphs (Additional file 1: Fig.S18). Although all topologies are spread widely across parameter space, different topologies are enriched in different regions of parameter space (Additional file 1: Fig.S6), and differ in their graph-size distributions (Fig.2(D), Additional file 1: Fig.S9). Particularly, acyclic graphs are more likely than cyclic graphs to be generated using gene regulatory rules where stable cell-types have smaller basins (Fig.2(C)). While the lineage graph topologies obtained at any parameter value do vary depending on *GR*, at the sample sizes used in this study, details of *GR* do not influence the distribution of lineage graph topologies (Additional file 1: Fig.S7)). To a large extent, these topologies can be characterized by their in-degree and out-degree distributions (Additional file 1: Fig.S8(A,B)). However, we find that acyclic graphs are slightly more enriched in our data than in randomized graphs with the same in-degree and out-degree distributions (Additional file 1: Fig.S8(C,D)).

#### Real homeostatic lineage graphs

Under normal circumstances, cellular differentiation is expected to be irreversible, therefore, we analysed the acyclic graphs of our model in more detail. We characterize acyclic graphs on the basis of two features: number of *branches* (*n_b_* = the total number of paths from root nodes to leaf nodes in the graph – 1), and number of *links* (*n_l_* = number of edges in the graph – number of edges in the maximal spanning tree of the graph) (Fig. 3(B), see also Additional file 1: Fig.S11). While for all chain type graphs, both *n_l_* and *n_b_* equal 0, for all tree type graphs, *n_l_* =0 and *n_b_* > 0. From Fig.3C, we see that the fraction of trees in our data remains low even if we relax the above definition of trees and allow *n_l_* to be non-zero. Even considering a threshold ‘treeness’ as high as *n_l_/n_b_* = 0.5, we find that the fraction of tree-type graphs in our data only increases from 0.01% to 0.013% (see also Additional file 1: Fig.S18). Therefore, in the following we use only the strict definition of trees, *n_l_/n_b_* = 0, as it does not affect our main conclusions.

**Figure 3.**
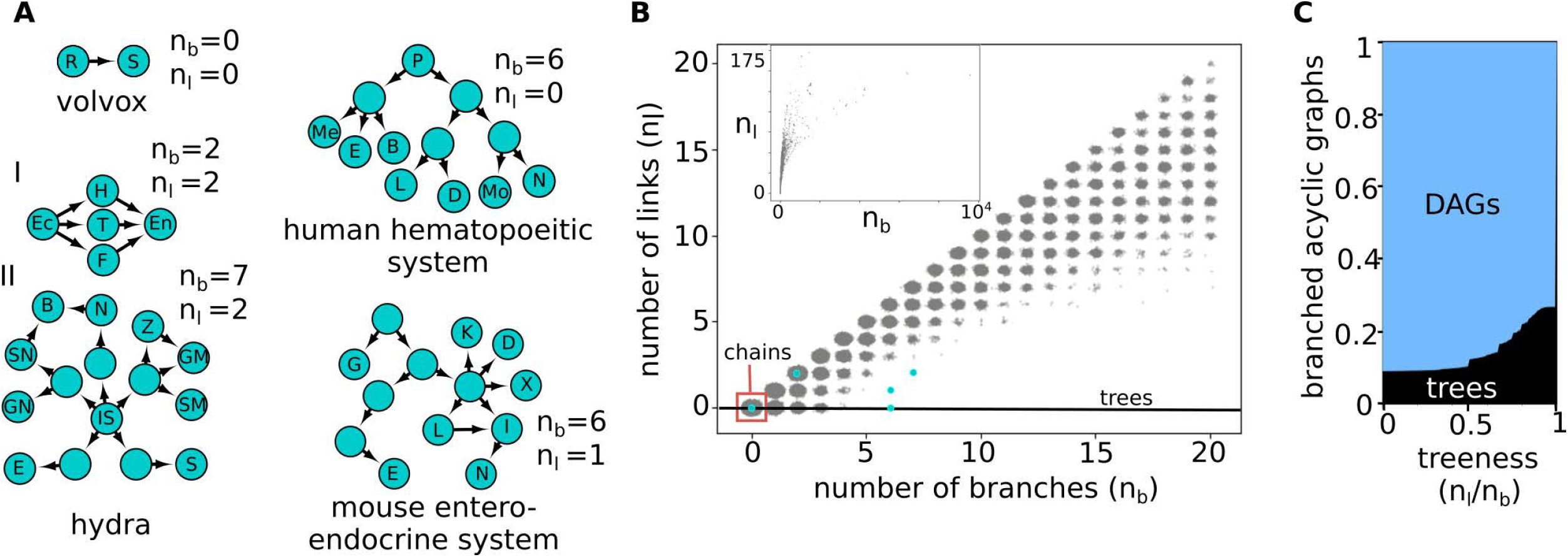
Lineage graphs of real organisms. **(A)** Lineage graphs from real organisms: Volvox: R= reproductive cell, S= somatic cell [33]; human hematopoietic system: P= progenitor cells, Me= megakaryocytes, E= erythrocytes, B= basophils, L= lymphocytes, D= dendritic cells, Mo= monocytes, N= neutrophils [41]; hydra: Ec= ectoderm, En= endoderm, IS = interstitial cell, H= hypostome, T= tentacle, F= foot, E= egg, S= sperm, GN= ganglion neuron, SN= sensory neuron, B= battery cell, N= nematocyst, Z= zymogen granule cell, GM= granular mucous granule cell, SM= spumous mucous granule cell [42]; and the mouse entero-endocrine system: G= goblet cell, E= EC cell, K= K-cell, D= δ-cell, X= X-cell, L= L-cell, I= I-cell, N= N-cell [43]. **(B)** Scatter plot of number of branches versus number of links for acyclic lineage graphs with *n_l_* and *n_b_* <= 20. The inset shows the *n_l_* versus *n_b_* scatter plot for all acyclic graphs. Noise has been added to points to make density of points more apparent. 1217108 graphs were used to generate this plot. **(C)** Relaxed definition of treeness: the x-axis represents *n_l_/n_b_*, our measure of a threshold for ‘treeness’. *n_l_/n_b_* = 0, represents the traditional, strict definition of trees, whereas at *n_l_*/*n_b_* = 1, all branched acyclic graphs are considered trees. At intermediate values, the fraction of graphs labeled as trees increases slowly. 521136 graphs were used to generate this plot.

We compare model-generated lineage graphs with examples of real lineage graphs collected from the literature (Fig.3(A)). While many reports of lineage graphs exist in the literature, especially owing to recent single cell transcriptomics studies, most graph reconstruction algorithms are biased to report trees [25, 26], and hence are not used in this study. We include only lineage graphs constructed in an unbiased way in this comparison.

The currently available real lineage graphs are still too few to statistically infer their features. Nevertheless, from this small sample it appears that real lineage graphs, especially mammalian ones, contain more branches and fewer links than model-generated graphs. This could indicate that the model is not sufficient to fully capture lineage graph topologies, or alternatively, that additional, unbiased real lineage graph reconstructions are required for rigorous analysis.

#### Homeostatic organisms contain pluripotent cells

We next looked at functional properties of model-generated homeostatic organisms; in particular we tested whether these organisms can regenerate using pluripotent cells. We define a pluripotent cell as any single cell-type which develops into a homeostatic organism using the same rules (GR, CD, *A* and SG) used to generate this organism in our data. We find that in 92.6% of organisms with acyclic lineage graphs (and 97% of all organisms) there is at least one pluripotent cell-type which is a part of the homeostatic organism. Since homeostatic organisms are stable products of the process of development, we consider them to be *adult organisms*. And we call pluripotent cells which are part of homeostatic organisms *adult pluripotent cells*.

Among real organisms, there exist both organisms which contain adult pluripotent cells (e.g. planaria), and those that do not (e.g. humans). In our data, in 82.9% of acyclic lineage graphs (73.3% of all graphs), pluripotent cells are more likely to be part the adult organism than not (Figs.4(A)).

**Figure 4.**
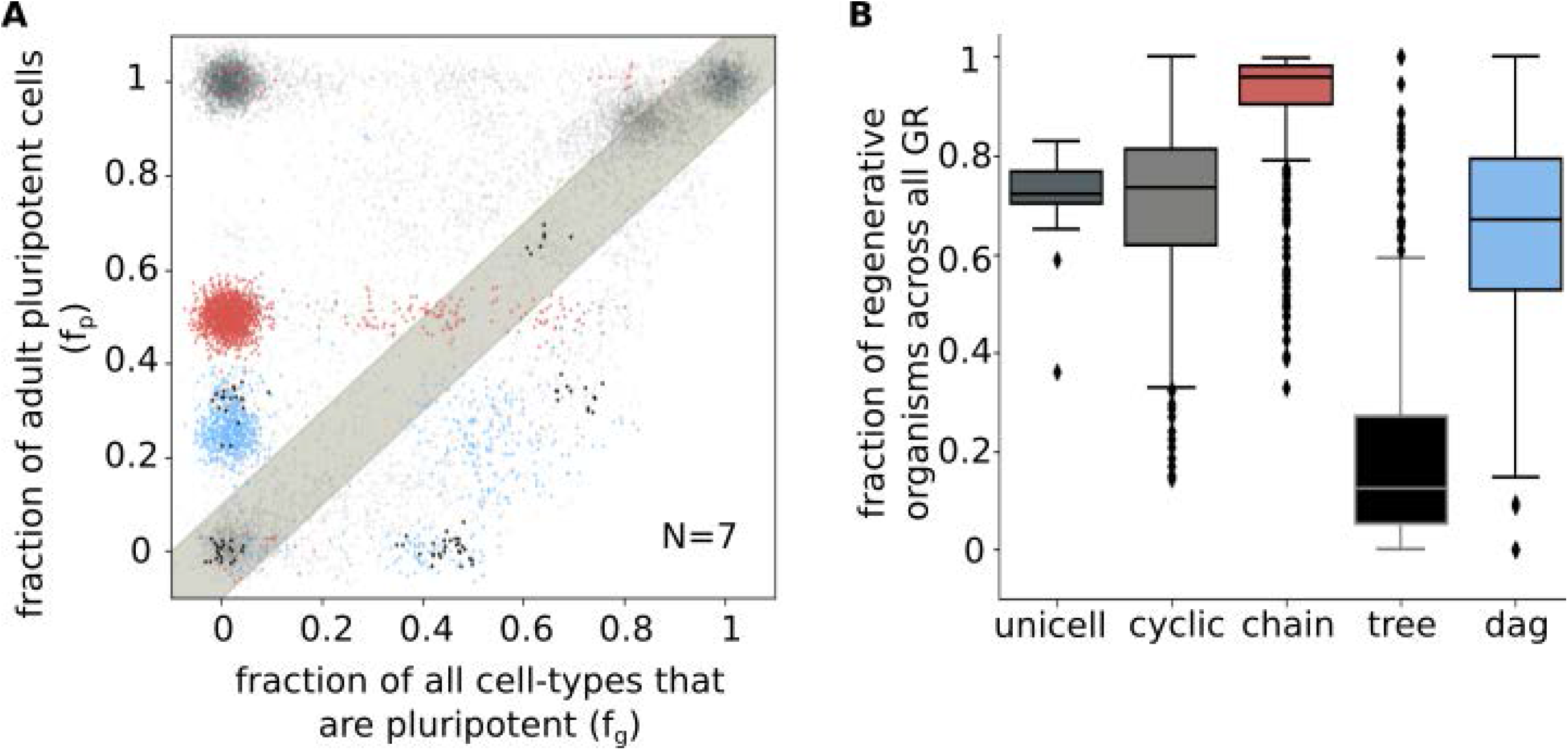
Regenerative capacity. **(A)** Scatter plot showing regenerative capacity for all *N* = 7 organisms generated with a fixed gene-regulation matrix GR using different matrices CD, *A* and SG (for cell-division, cellular adjacency and signal transduction, respectively). Each point represents an organism. The x-axis is the fraction of all cell-types that are pluripotent (*f_g_*). The y-axis is the fraction of cell-types in the organism that are *adult pluripotent* cells (*f_p_*). Noise has been added to the position of points to make their density more apparent. Colours of points indicate the topology of their lineage graphs (as in Fig.2): unicellular:dark grey, cyclic:light grey, chain:red, tree:black, DAG:blue. Points above the grey band are regenerative organisms (with *f_p_ / f_g_* > 1). 13177 graphs were used to generate this plot (see also Additional file 1: Fig.S12). **(B)** Box plot of proportion of regenerative graphs of different topologies across all organisms in the data (see also Additional file 1: Fig.S13(A)). For each GR used in our data, for a given graph topology, we looked at the fraction of graphs with regenerative capacity > 1 (equivalent to the fraction of points of a certain colour that occur above the grey band in **A**). Boxes represent quartiles of the data set. Lines inside the box show the median, while whiskers show the rest of the distribution. Outliers are shown as diamonds. 4852994 graphs were used to generate this plot.

Evidently, organisms that contain adult pluripotent cells are more disposed to be regenerative than those that do not. We therefore measure *regenerative capacity* of an organism as the fraction of cell-types in the organism that are adult pluripotent cells (*f_p_*) divided by the fraction of all cell-types (irrespective of whether it is a part of the adult organism, or not) that are pluripotent (*f_g_*). We call an organism *regenerative* if its regenerative capacity is greater than 1. We find that regenerative capacity differs among different topologies (Fig.4(B), Additional file 1: Figs.S13(A)). Notably, while most chains are regenerative, most tree-type graphs have very low regenerative capacity (see Additional file 1: Fig.S19 for regenerative capacities using relaxed definition of trees). More generally, this distribution of regenerative capacities serves as an example of correlations between high-level functions of organisms with their lineage graph topologies. Such associations could form the basis for natural selection favoring certain topologies over others in real multicellular organisms.

#### Signal transduction influences regeneration trajectories

The high level of adult pluripotent cells in homeostatic organisms is surprising, since cell fates within the organism are constrained due to signaling, and cells taken out of the context of signaling from other cell-types in the organism, are not expected to regenerate the other cell-types.

In order to test the mechanism of regeneration in the model, we asked two questions: 1) how much does cell-fate in the organisms depend on signaling, and 2) what do regeneration trajectories look like? If regeneration does not depend on signaling, cell-types should exactly retrace their paths in the homeostatic lineage graph, irrespective of the presence of other cell-types (Fig.5(A)).

**Figure 5.**
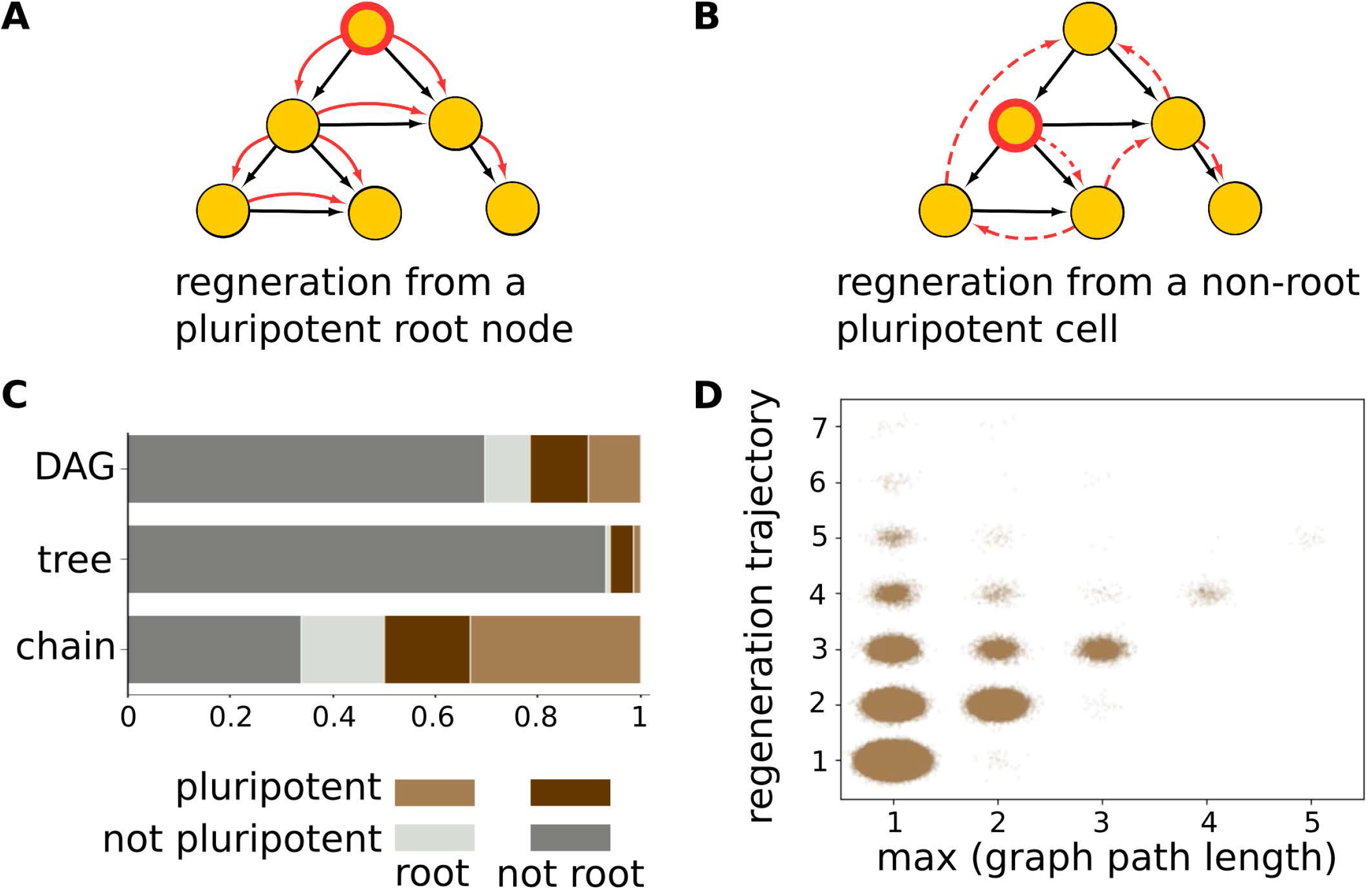
Regeneration trajectories. **(A,B)** Schematics of regeneration trajectories.Yellow circles represent cell-types in a homeostatic organism, and black edges represent lineage relationships. Adult pluripotent cells are outlined in red. Red edges represent lineage relationships between the organism’s cell-types during regeneration. **(A)** Here the root node is the only adult pluripotent cell and regeneration trajectories exactly match the paths in the lineage graph. In cases where signaling does not play a role in determining cell-fate, all regeneration trajectories are of this kind. **(B)** Here, a non-root node is a pluripotent cell, and therefore necessarily, regeneration trajectories cannot be perfectly aligned with paths in the lineage graph. In these cases, signaling is definitely involved in governing cell-fates. Dashed red edges imply that cell-types other than those present in the homeostatic organism may be produced during regeneration. **(C)** Stacked histograms for cell-types of different categories pooled from organisms with different lineage graph topologies. Different cell-type categories are represented with different colours. Non-pluripotent cells are represented in greys; root nodes: light grey, not non-root nodes: dark grey. Adult pluripotent cells are represented in colours; root nodes: light brown, non-root nodes: dark brown. Heights of colored blocks represent the proportions of corresponding cell-types. 1217108 graphs were used to generate this plot. **(D)** Scatter plot showing maximum path lengths across lineage graphs vs lengths of regeneration trajectories from pluripotent root nodes for acyclic lineage graphs that contain them. Each point represents an organism. Noise and transparency has been added to the position of points to make their density more apparent. 699986 graphs were used to generate this plot.

To answer the first question, we define the fate of a cell-type *C* in a homeostatic organism as the set of all cell-types that receive an edge from cell-type *C* in the lineage graph of the organism. Note that the above definition refers to a *proximal* cell-fate, where only the immediate descendants of a cell-type are considered. We call a cell-type *independent* if it has the same cell-fate when taken out of the homeostatic organism, as it does within the organism. We find that across all parameter regions, homeostatic organisms are enriched in independent cell-types (Additional file 1: Fig.S14). While about 32% of all independent cells, pooled from all acyclic graphs, are adult pluripotent cells, 90% of all adult pluripotent cells are independent (Additional file 1: Fig.S15). That is, in the model, proximal cell-fates, especially those of adult pluripotent cells, are likely to be independent of signaling.

To resolve the second question, note that if regeneration trajectories recapitulate the homeostatic lineage graph, (a) only *root nodes*, which are nodes from which all other nodes of the lineage graph are reachable, can be adult pluripotent cells, and (b) regeneration trajectory lengths must match the longest of the minimum path-lengths from the root node to any of the leaf nodes in the lineage graph.

Now, necessarily, there can be no more than a single root node in an acyclic graph. Some acyclic graphs, for example convergent trees (Additional file 1: Fig.S10), even lack root nodes. In our data, only 7.4% acyclic graphs lack a root node, and of these, 85.4% indeed lack adult pluripotent cells. And, 90.4% of the time, in rooted acyclic graphs, lengths of regeneration trajectories starting from pluripotent root nodes do match the maximum path length in the respective lineage graphs (Fig.5(D)). But, among these rooted, regenerative acyclic graphs, only 62.9% have root nodes that are pluripotent. Overall, 41.6% of adult pluripotent cells are *not* root nodes (Fig.5(C)). Regeneration trajectories starting from such non-root adult pluripotent cells are bound to be different from paths in the homeostatic lineage graph (Fig.5(B)), and therefore must involve signaling. Consistently, lineage graphs with non-root adult pluripotent cells are more likely to be generated at higher values of *P*_sig_ and *P*_adj_ than lineage graphs with only root pluripotent cells (Additional file 1: Fig.S16).

To summarize, although cell-fates for most cell-types in organisms are independent of signaling, almost half the time, signal exchange definitely plays a role in regeneration, and regeneration trajectories of organisms are different from paths in their homeostatic lineage graphs. Parallels to these results can be seen in the development and regeneration of ascidians, which undergo ‘mosaic development’, where cell fates are independent of cellular context, and are determined by *autonomous specification* [27]. But the level of plasticity during asexual budding (which can be seen as a form of programmed whole body regeneration) in colonial ascidians, which are model organisms both for mosaic development and regeneration, points to a ‘non-mosaic’ mode of regeneration [28].

## Discussion

The process of development and its molecular mechanism are inherent in all *metazoans* and in all plants [3]. It is therefore difficult to design experiments that could distinguish between emergent traits associated with development and traits that have evolved on top of it. Here, we have developed a minimal model where we can look at development in the absence of complications due to cross-talk with other biological processes. In our model, we include only those ingredients of development that are shared across all multicellular organisms, while not ascribing any particular form or mechanism to these processes. This allows us to identify universal traits that are inherent to development, regardless of the details of the process, or the distinct selective pressures different organisms may be subject to. We see such a prominent emergent trait in our model: ability of *whole body regeneration* (WBR) through pluripotency. Concurrently, WBR, although absent in many animals, such as mammals, is widely spread across basal metazoan phyla.

Note that such basic traits can still be subject to selection through regulatory processes on top of the key ingredients of development. Below, we discuss major assumptions and limitations of our model, and contrast these with mechanisms that occur in biological organisms.

- *Considerations of space and time:* Spatial arrangement of cells and cell movements are important and highly regulated aspects of development. Two cells having the same cell-type but occupying different niches are likely to receive different sets of signals, and therefore are expected to behave differently [12]. Cells are also conjectured to lose potency as development progresses, such that cell-types present in later stages of development are likely to give rise to fewer cell-types than those present in the earlier stages [29]. Such a process can be captured in the model by using developmental rules endowed with a *flow* such that cell types appearing later in development have lower potency. The final outcome of development is highly likely to be affected in those organisms where such a mechanism operates. Although in the current study, due to our focus on widely exploring asymmetric cell division, signaling and gene regulation, we do not explore the important aspects of space and time, we provide a recipe for how they can be included in future studies: in Additional file 1: section 1.1, using the example of the *Drosophila* segment polarity network, we show how space can be encoded within the framework of the model. Developmental time can be included using a similar approach.
- *Independent processes:* Cell-division, signaling, and gene regulation are treated as independent processes in the model. This is likely to be false in real animals. Primarily, this implies that not all regions of parameter space explored in this work are biologically feasible. In particular, cells in the model follow a simple program for asymmetric cell division that is intrinsic to cell-types. But extrinsic control of asymmetric cell division, involving cues from surrounding cells, does occur in animals [18]. Extrinsic control of asymmetric cell-division could lead to a decrease in the independence of cell-fates on cellular context which we see in the model, which could affect regenerative capacity.
- *Chemical signaling:* In our model, signal recognition is based on identities of the donor and the recipient cells. In contrast, in real organisms, cells contain receptors that recognize signal molecules, rather than recognizing the donor cells that produce those signals. Firstly, since there are fewer kinds of signal molecules than there are cell-types, it is likely in this chemical recognition scheme that a cell-type will receive the same set of signals even if some other cell-types in the organism are changed. That is, cell-fate is likely to be even more robust to changes in cellular context than the present model. In our model, cell-fates of most pluripotent cells are independent of cellular context. Therefore, a version of the model with chemical recognition is also likely to yield regenerative organisms.
- *Other schemes:* In the current model, we use the following scheme of development: cell-division, followed by signaling among daughter cells and gene regulation in response to signals exchanged. But there are other reasonable schemes which can also be considered. For example, a scheme where cell-division is followed by an additional step of gene regulation before signal exchange is also plausible. In the current model, daughter cells contain subsets of the contents of the mother cell, and in this sense are more similar to each other than to daughters of other mother-cells. Therefore, in the current scheme, signals received from a sister cell are likely to be less *effective* in changing cell-state than are signals received from other daughter cells. Gene regulation right after cell division would lead to a diversification of daughter cells, which is therefore likely to increase the level of effective signaling among daughter cells. In our model, level of signal exchange is controlled by *P*_adj_, and we find that pluripotency increases, albeit modestly, with *P*_adj_ (Additional file 1: Fig.S13(E)). Therefore, we expect that switching to this other, more elaborate, scheme of development would still lead to high regenerative capacity.
- *Additional parameters:* The effect of processes such as asynchronous gene state updates [30], and time delays involved in transfer of information about gene state updates [31] has been tested on the *Drosophila* segement polarity network, and found to have interesting effects on the robustness of phenotypes. Such processes could add to the richness of lineage graphs we obtain from our model, but come with the cost of additional parameters, which would limit the breadth of the sampling.
- *Cell death:* Cells in the model do not die. Not including cell death in the model results in lineage graphs where each node has at least one out-edge. We anticipate that including cell death would reduce the number of cycles in lineage graphs, leading to an increase in the proportion of acyclic graphs (Additional file 1: Fig.S17). Since regenerative capacity is linked to lineage graph topology, cell-death could be an important factor in determining regenerative capacity. We also anticipate that including cell death could add a sense of developmental time to the model.

Although here we only provide intuitive arguments for what alternate versions of the model might yield, the framework of the model is easily amenable to manipulations, and differently constructed versions can be tested in the future.

The present model makes several predictions regarding general features of development and multicellular organisms. It suggests that presence of *adult pluripotent cells* should be a widespread trait in multicellular life-forms. In plants, we are already aware of pluripotent cells in the root and shoot meristems. But among animals, a wider investigation of regeneration and its mechanisms will be required to test this idea. A recent example of such a study is [32], where the authors test the ancestral nature of regeneration in *Nemertaean* worms, which are not classical model organisms.

The distribution of acyclic lineage graph topologies in our data reflect the complexity and diversity of forms of multicellular animals that biological development is expected to produce. Small (2-node) *chains* are the most abundant acyclic lineage graphs in our data (Fig.2(D), Additional file 1: FigS9(C)). In line with this, the simplest multicellular organisms, such as *Volvox carteri* [33], an alga which evolved multicellularity only recently, has a chain like lineage graph. Interestingly, some cyanobacteria, such as *Anabaena spaerica* [34], which display multicellularity during nitrogen starvation, also have chain-like lineage graphs.

*Tree-type lineage graphs are rare* in our data (Fig.2(A,B)), and tend to be small, and convergent rather than divergent (Additional file 1: Figs.S9(E), S10). This could indicate one of two things: This could imply that lineage graphs of complex organisms are unlikely to be tree-like. Our data suggests that they are more likely to be DAGs (directed acyclic graphs), i.e., organisms have higher levels of trans-differentiation than expected (Fig. 3(B), Additional file 1: Fig.S11). Or, it could mean that more complex regulation, on top of the ingredients of this model, are at play in real organisms which lead to complex tree-like lineage graphs. A perhaps presumptuous, but interesting possibility is that tree-like lineage graphs were selected for because of their low regenerative capacity. There exist arguments and speculation over whether mammals, among other animals, selectively lost the ability to regenerate, and why [35].

These questions surrounding the topologies of lineage graphs are likely to be resolved very soon in the future, given the rapid developments in single cell transcriptomics technology. A notable recent study is that of Plass *et al*. [36], where they assemble the whole organism lineage graph for *Planaria*. A possible hurdle comes from the fact that in such studies, lineage relationships cannot be directly accessed and are instead inferred using distances between cellular transcriptomes obtained at different times. Moreover, current methods for lineage reconstruction using single cell transcriptomics data are not unbiased; in [36], although lineage reconstruction yielded a complex graph, the authors highlight the best supported spanning tree of this graph. Current lineage reconstruction methods work best if a particular topology for lineage graphs is already anticipated, and most methods are designed to only find chains and trees[26, 25]. In contrast, a study by Wagner *et al.* [37], where single cell transcriptomics is used in conjunction with cellular barcoding, provides an example of a lineage reconstruction method which is unbiased towards particular topologies. In agreement with our result, the authors of this study found that zebrafish development is best represented by a DAG.

Lastly, we discuss how certain predictions of our work can be experimentally tested. Our work suggests that in colonial ascidians, which reproduce asexually using a variety of budding structures, the pattern of regeneration should be ‘non-mosaic’; where regeneration trajectories do not recapitulate lineage trajectories in the homeostatic organisms. In contrast, our model suggests that in *Planaria*, where pluripotent c-Neoblasts appear to occupy the root node [36], regeneration trajectories are likely to reflect the homeostatic lineage graph. These predictions can be addressed by lineage reconstruction experiments that compare homeostatic lineage graphs with lineage graphs produced during regeneration of these organisms.

Our results also suggest that in organisms such as *Planaria* and perhaps colonial *Ascidia*, where regeneration is based on adult pluripotent cells [15, 16], these cells are likely to be independent of cellular context; that is, their proximal cell fates should not change when taken out of the body, or transplanted to other cellular contexts. In *Planaria*, c-Neoblast independence could explain the coarse pattern of distribution of specialized neoblasts across the planarian body, and also why the distribution of specialized neoblasts produced does not depend on which organ is amputated [15]. Recent development of a method to culture neoblasts in the lab [38], make it possible to experimentally test neoblast independence.

## Conclusions

Development transforms single-celled zygotes into multicellular adults by combining three basic processes: asymmetric cell-division, cell-signaling, and gene regulation. Despite undergoing this common set of developmental processes, multicellular organisms display a huge variety of forms. In this work, we use a generative model of development to gauge the extent of possible diversity of multicellular forms. We explore the forms of ‘organisms’ generated by our model in terms of their cell-type lineage maps. Our data indicates that cell-type lineage maps are unlikely to be tree-like, and instead that organisms are likely to undergo a much higher level of trans-differentiation than anticipated. Additionally, ‘organisms’ generated in our model contain adult pluripotent cells, and are thus likely to be capable of whole body regeneration. This observation supports the view that whole body regeneration is an epiphenomenon of development. Regenerative capacity differs among organisms with different lineage graph topologies, tree-type lineage graphs having the lowest regenerative capacities. These differences could potentially serve as a basis for selection biased towards certain topologies. Our results also suggest that regeneration trajectories are likely to deviate from paths in the cell-lineage graph, and could involve cell-types not present in the adult organism. To summarize, our work ties together the process of development and the phenomenon of regeneration, and suggests many testable hypotheses, which can be addressed through experiments on well-established model organisms, but more importantly, through a wider sampling of cell-differentiation trajectories across multicellular organisms.

## Methods

### Surveying the combinatorial space of developmental schemes

We considered organisms with *N* = {3, 4, 5, 6, 7} genes. For each *N*, we have looked at {100, 100, 100, 92, 25} randomly generated gene regulation matrices (GR), respectively. For each GR, all values from [0, 0.1, 0.2, …, 1.0] were used for the parameters *P*_asym_, *P*_sig_ and *P*_adj_. A distinct set of developmental rules matrices (CD, *A* and SG) were generated for each set of parameter values. And for each set of rules matrices, 10 randomly chosen cell-types were used as initial conditions (in case of *N* = 3, all 8 cell-types were used). In all, we have looked at about ((100 + 100 + 100 + 92 + 25) × 11^3^ × 10) ≈ 5.5 × 10^6^ systems. 4858643 of these converged within 1000 time-steps into homeostatic organisms.

### Model details

All codes used to generate and analyse data are written in Python3.6 or Octave 5.2.0.

#### Asymmetric cell division

In our model, for any cell-type *C_i_*, we generate different sets of daughter cell-types *D_i_* using the parameter *P*_asym_ ∈ [0,1]; for any daughter cell-type *D*_*i*_*i*1__ ∈ *D_i_*, ∀*k* ≤ *N*,

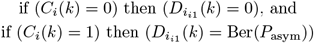

We encode cell-division in a binary matrix CD_2*^N^* × 2*^N^*_; CD(*i, j*) = 1 if cell-type *C_j_* ∈ *D_j_*, else CD(*i, j*) = 0 (Fig.1(B)).

#### Signaling

The probability that a gene in the model produces a signaling molecule is *P*_sig_ ∈ [0,1]. Formally, let SG = {0,1}*^N^* be a binary vector. Then gene *k* produces a signaling molecule if SG(*k*) = 1, where SG(*k*) = Ber(*P*_sig_) (Fig.1(C)). Let SG*_j_* = {0,1}^*N*^ be the set of signals produced by cell-type *C_j_*. For any gene *k*, SG*_j_*(*k*) = 1 ⟺ (*C_j_*(*k*) = 1) ∧ (SG(*k*) = 1).

Parameter *P*_adj_ ∈ [0,1] gives the probability of signal reception. We encode signal reception in a binary matrix A_2^*N*^_ × _2^*N*^_. Cell-type *C_i_* receives all signals produced by cell-type *C_j_* if *A*(*j, i*) = 1, where *A*(*j, i*) = Ber(*P*_adj_). *C_i_* receives no signals from cell-type *C_j_* if *A*(*j, i*) = 0 (Fig.1(D)). Cells can only receive signals from other cell-types present in the same time step. Let *T_t_* = {0,1}^2^*N*^^ be a binary vector, where *T_t_*(*i*) = 1 if cell-type *C_i_* is present in the time step *t*. *T_t_* represents the state of the organism at time step *t*. For some cell-type *C_i_* present at time step *t*, let 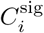 represent its state immediately after signal exchange. In cell-types that receive a signal, the corresponding genes are set to 1 (Fig.1(F)). That is,

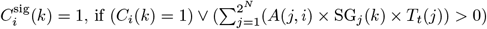

#### Gene regulation

We define gene regulation in the model as a set of stable cell types, and cell-types in the basins of these stable cell-types. As mentioned earlier, stable cell types need not be fixed points (single cell-state) of the gene regulatory network, they can also be an oscillation (multiple cell-states). Oscillatory stable cell-types are represented as the set of all cell-states that compose the oscillation.

Formally, a system with *N* genes can have *n* ≤ 2*^N^* stable cell-types {*S*_1_, *S*_2_, …,*S_n_*}; where *S_x_* is itself a collection of *n_x_* cell-states {*C*_*x*_1__, …*C*x_n_x__** } such that *x*_1_ < *x*_2_ < … < *x_n_x__*. For any two cell-types *S_x_* and *S_y_*, if *x* < *y* then *x*_1_ < *y*_1_.

We encode gene regulation in a binary matrix *GR*_2^*N*^_ × _2^*N*^_. To generate *GR* for a given organism, we pick the number of stable cell-types *n* ≤ 2^*N*^ according to uniform random distribution. First, we assign cell-states that form the basins of these stable cell-types: Cell-states are uniform randomly partitioned among the *n* basins. We then choose cell-states that form the stable cell-type from within the corresponding basins: let *B_x_* be a basin, then for some *j* such that (*C_j_* ∈ *B_x_*), (*C_j_* ∈ *S_x_*) with probability 0.5. For all *i* such that *C_i_* ∈ *B_x_*, GR(*i, j*) = 1 if (*C_j_* ∈ *S_x_*).

#### Homeostatic organisms and their cell-type lineage graphs

Let us consider an organism in state *T_t_* at time step *t*. Right after cell division, let the state of the organism be represented by 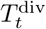. After division, the organism is composed of all the daughter cells produced in that time step. That is,

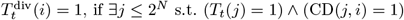

These daughter cells exchange signals among themselves. Let 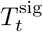 represent the state of the organism right after signal exchange. Then,

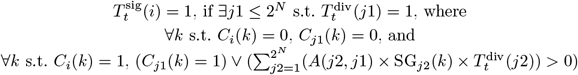

The signals received by a cell-type activates its gene regulatory network. Gene regulation updates the set of cell-types according to the following expression: ∀*i* ≤ 2^*N*^,

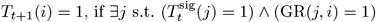

Therefore, the organism is only composed of stable cell-types. Let the system have *n* ≤ 2^*N*^ stable cell states. Then, we can equivalently represent the state of the organism at time step *t* as a binary vector 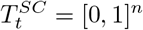, such that for *x* ∈ {1, 2, …*n*}.

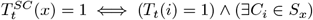

We call states of the organism such that 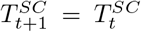 homeostatic organisms (Fig.1(F,G)).

We represent the homeostatic organism as a cell-type lineage graph. The nodes of the graph represent stable cell states that are present in the homeostatic organism, and directed edges represent lineage relationships between these stable cell states. Let the stable cell states *S*_*x*1_ and *S*_*x*2_ both be present in the final organism, and let them be represented by nodes *V_a_* and *V_b_* of the lineage graph respectively. Then, there is an edge from *V_a_* to *V_b_* if one of the daughter cells of *S*_*x*1_ gives rise to *S*_*x*2_ after one round of cell-signaling and gene regulation (Fig.1(G)). That is,

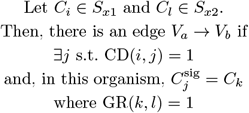

### Assignment of topologies to lineage graphs

We categorize lineage graphs into 6 topologies: unicellular, strongly connected component(SCC), cyclic, chain, tree and other directed acyclic graphs (DAG). We ignore self-edges while assigning these topologies. A lineage graph is called *unicellular* if it has only a single node. For all other topologies, we used the networkx (version 2.2) module of Python3.6. A lineage graph is called *SCC* if the graph has more than 1 node and contains a single strongly connected component, it is called *cyclic* if the graph contains cycles and has more than one strongly connected component, it is called a *chain* if networkx classifies it as a tree and the maximum in-degree and out-degree are 1, it is called a *tree* if networkx classifies it as a tree and maximum in-dergee or out-degree is greater than 1, and it is called a *DAG* if networkx classifies it as a directed acyclic graph but not a tree.

### Lineage graph randomization protocol

We represent a lineage graph with *e* edges as a matrix *E*_*e*×2_, where *E*(*i*, 1) and *E*(*i*, 2) represent the source and the target node of edge *i* respectively. To randomize lineage graphs, we used a protocol that preserves in and out degrees of each node; we randomly choose pairs of edges from the graph and swap their target nodes. Let the randomized graph *E*_rand_ be initially identical to *E*. Then,

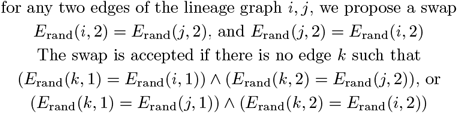

The above condition ensures that the total number of unique edges in *E* and *E*_rand_ remain the same. We swap edges 1000 times for each lineage graph to randomize it.

### Independent and intrinsically independent cell-types

We call a cell-type *independent* if it has the same cell fate when grown outside the organism as it does when it is a part of the organism. The cell fate 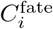 of some cell-type *C_i_* in the organism is given by the set of cell-types receiving an edge from the node *C_i_* in the organism’s lineage graph. To decide whether a given cell-type *C_i_* is independent or not, we separate this cell-type from the rest of the organism, and allow it to undergo one round of cell division, signaling and gene regulation, according to the same matrices CD, SG, A and GR that were used to generate the organism from which it was taken. Let us call the resulting set of cell-types 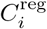. We call the cell *C_i_* independent if 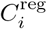 is identical to 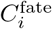.

For some cell-types, the basis of their independence is an insensitivity to signals produced in the organism. In such a case, the set of signals produced by the daughter cells of the cell-type is sufficient to satisfy the maximum set of signals that each of the daughter cells can receive.

Let the set of daughter cells of cell-type *C_i_* in an organism be *D_i_*. ∀*C_j_* ∈ *D_i_* let 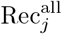 represent the maximal set of signals that it can receive, when all 2^*N*^ possible cell-types are present together. i.e., For all signaling molecules k such that SG(*k*) = 1,

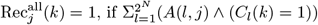

And, let 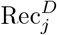 be the set of signals it receives from within the set of cells *D_i_*. i.e.,

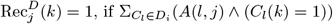

If, for all *C_j_* ∈ *D_i_*, 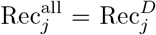, *C_i_* is *intrinsically independent*.

## Acknowledgements

We thank Luca Peliti, Albert Libchaber, Mukund Thattai and John McBride for useful discussions, and John McBride for assistance with writing Python code for analysis.

## Funding

This work was supported by the taxpayers of South Korea through the Institute for Basic Science, Project Code IBS-R020-D1.

## Abbreviations

DAG: Directed Acyclic graph
RBN: Random Boolean Network
ER: Erdos-Renyi
SCC: Strongly Connected Component
WBR: Whole Body Regeneration

## Availability of data and materials

Octave5.2.0 and Python3.6 code were used in this work. All data generated or analysed during this study are included in this published article and its supplementary information files. All data, and codes used to generate and analyse the data, are available in BioModels repository [39] and assigned the identifier MODEL2103180001 [40].

## Ethics approval and consent to participate

Not applicable

## Competing interests

The authors declare that they have no competing interests.

## Consent for publication

Not applicable

## Authors’ contributions

S.M. conceived the project, developed code for simulations and performed analysis; S.M. and T.T. designed research; S.M. and T.T. wrote the paper. All authors read and approved the final manuscript.

## Author details

Living Matter Theory Group, Institute of Basic Science-Center for Soft and Living Matter, Ulsan, South Korea.

## 1 Supplementary material

### 1.1 Drosophila segement polarity network expressed in terms of the generative model

In Drosophila embryos, segment polarity genes maintain borders of parasegments, which are 4 cells wide. Within each parasegment, the polarity genes are expressed in characteristic stripes. In Albert and Othmer (2003), the authors demonstrated that the gene regulatory network responsible for the pattern of gene expression in this system can be modeled as a Boolean logical network. In the following, we examine the Drosophila segment polarity network in terms of our generative model.

The network consists of 15 nodes: en, wg, hh, ptc and ci represent mRNAs, and SLP, EN, WG, HH, PTC, SMO, PH, CI, CIA and CIR represent proteins. Of these, WG, hh and HH act as signals. Signaling molecules HH and WG do not participate in regulation within the cells that produce them, rather they act only in cells that receive them as signals. In order to incorporate this feature, we represent each cell in the parasegment as two model cells; production of all non-signal molecules takes place in one of the cells, and molecules responsible for regulation of signal molecule production are exported to the second cell, from which signal molecules are secreted (Fig.S1(A,B)). In this sense, the second cell acts as a special compartment which insulates the gene network in the first cell from regulation by signal molecules produced within the same cell.

In this system, signals are exchanged only between neighbouring cells. Accordingly, in our model, cell positions can be expressed as additional ‘genes’, whose states do not change. For example, to express the positions of the 4 × 2 cells in this system, we use 3 additional ‘genes’ (Fig.S1(C)). In this system, signal exchange only depends on these ‘positional genes’, and does not depend on the states of the other genes.

Thus, our model is capable of expressing spatial arrangement of cells, and complex signaling mechanisms, although, it comes at the cost of an increase in system size. In Fig. S2, we show the signaling vector SG, and portions of the cellular adjacency matrix *A*, and gene regulation matrix *GR* relevant to the steady state of the wild-type segment polarity network. The authors assume symmetric cell-division in Albert and Othmer (2003), and we do the same; therefore we do not show the cell-division matrix CD here.

**Figure S1:**
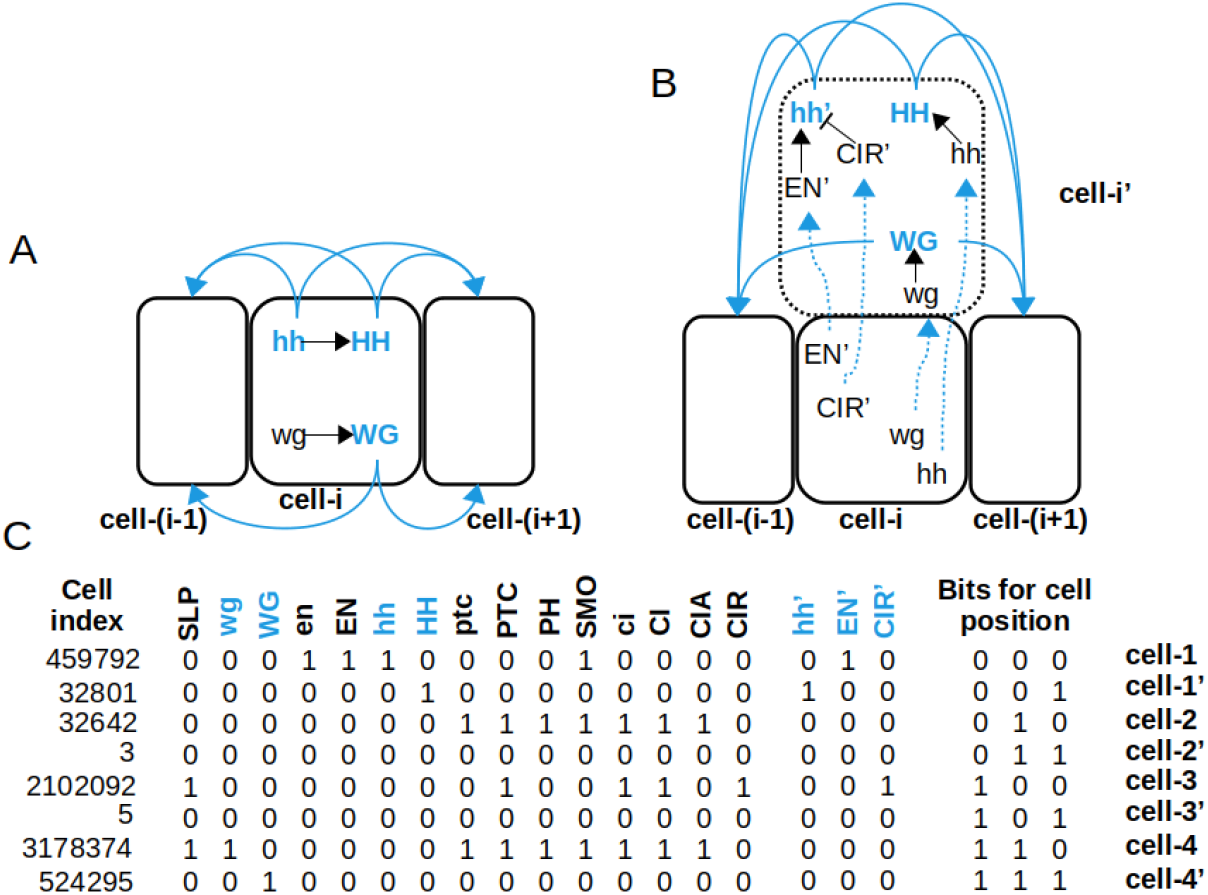
Modified segment polarity network. **(A)** Signaling in the original model, as in Albert and Othmer (2003)). Signaling molecules are labeled in blue. Blue edges represent signal transduction, and black edges represent ‘gene’-regulation. **(B)** Modified structure of segment polarity network. We introduce a new cell, shown here with a dotted outline, adjacent to the original cell, which acts like an insulated compartment of this cell. All signals are transmitted via this new cell to neighbouring cells. **(C)** Steady state of the modified network that corresponds to wild-type stripe pattern in Drosophila, as reported in Albert and Othmer (2003). Each row is a cell-type, and columns represent states of ‘genes’. 1 implies presence of the gene product, and 0 implies absence of the gene product. There are 21 genes in the system: the first 15 genes are the original mRNAs and proteins used to construct the regulatory network in Albert and Othmer (2003), and the next 3 genes represent ‘mirrors’ of hh, EN and CIR which are used for signaling purposes. The last 3 ‘genes’ encode the position of the cell along the anterio-posterior axis; cell-1 is the most anterior and cell-4 is the most posterior. Cells 1’ −4’ represent the new cells we introduce for signal transduction. Each cell is indexed by the decimal number obtained upon converting the corresponding 21-length binary vector.

**Figure S2:**
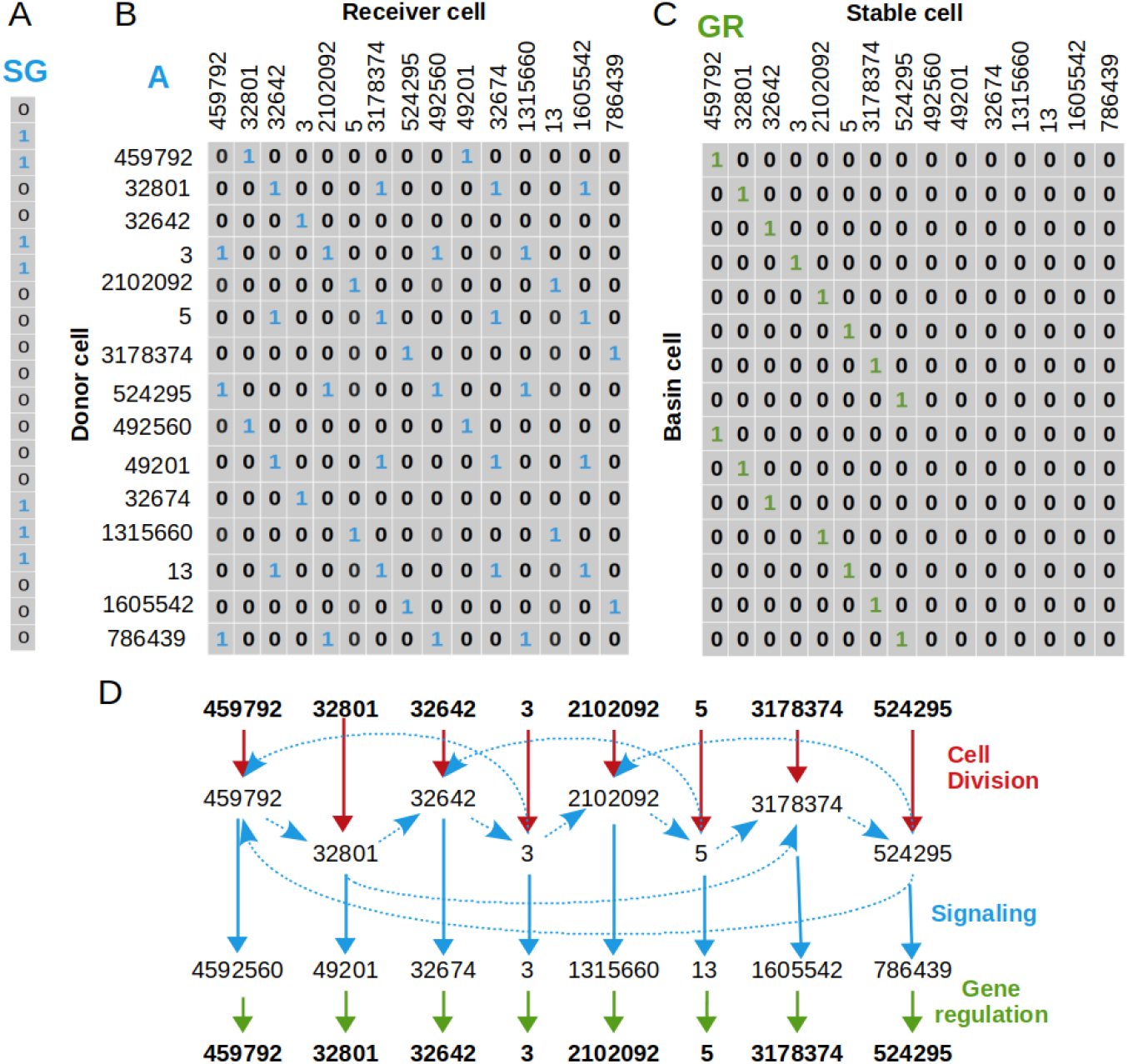
Developmental rules matrices for the modified segment polarity network. **(A)** signaling vector SG, **(B)** Cellular adjacency matrix *A*. Note that cellular adjacency is completely determined by cell positions; Cells-1-4 only pass on molecules to cells-1’ −4’ respectively, and a cell-i’ only passes on signals to cell-(i-1) and cell-(i+1). Periodic boundary conditions are employed here, which implies that cell-1 and cell-4 are neighbours. **(C)** Gene regulation matrix *GR*. The first 8 cell-types correspond to the stable state, as in Fig.S1(C). In **B** and **C**, only the relevant parts of the rules matrices are shown. The full matrices are of size 2^21^ × 2^21^. **(D)** Schematic diagram of signaling and gene regulation in determining the wild-type steady state of the *Drosophila* segment polarity network. Numbers represent the indices of different cell-types. Red arrows represent cell-division, which is symmetric in this case. Dashed blue arrows represent signal exchange among cell-types and solid blue arrows represent changes in cell-types due to signal exchange. Green arrows represent gene regulation.

### 1.2 basin sizes of gene regulatory networks

In the model, stable cell-types and cells in their basins of attraction are randomly chosen. In gene regulatory networks with N genes, between 50% - 75% of *2^N^* cells are stable cell-types (Fig.S3(B)). On average, basins of these stable cell-types are small, and most basins contain a single cell-type (Fig.S3(B,C)).

**Figure S3:**
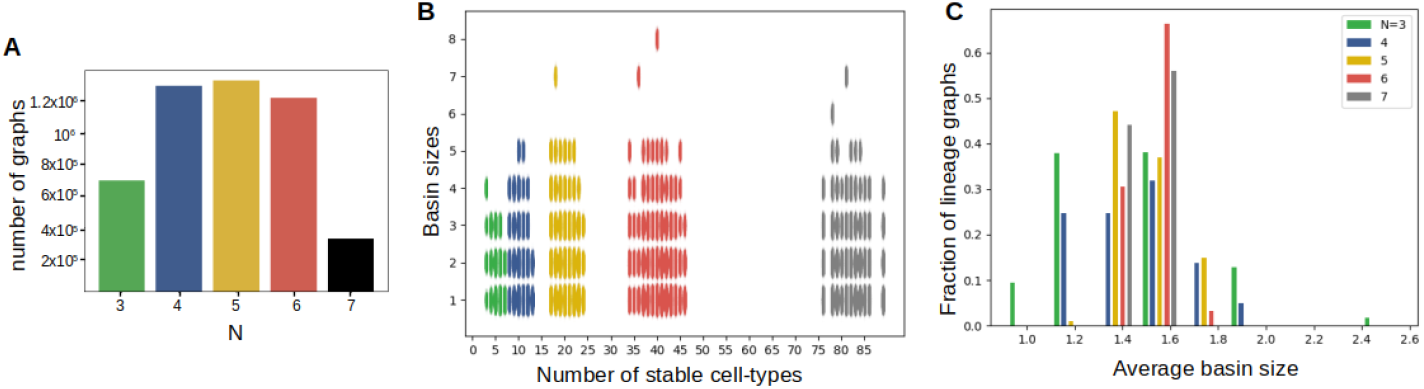
Distribution of basin sizes of lineage graphs. 4852994 graphs were used to generate these plots. **(A)** Number of model generated homeostatic organisms in the data at different values of N. **(B)** Basin sizes in organisms containing different numbers of stable cell-types. (**C**) Histogram of average basin size for all homeostatic organisms.

### 1.3 Lineage graph size

While a majority of graphs in our data are small (1-5 nodes), the largest graphs have 89 nodes (Fig.S4(A)). The number of nodes in lineage graphs follows closely the diversity of daughter cell-types produced (Fig.S4(C,D)). At very low *P*_asym_, cells produce daughters cells identical to themselves, and at very high *P*_asym_, most daughter cells are of the type [0,0, …0]. Therefore at these values, diversity of daughter cells, and correspondingly the number of nodes in lineage graphs, is low. At other values of *P*_asym_, the number of nodes stays level and decreases slowly beyond *P*_asym_ = 0.5 (Fig.S4(C)). Number of nodes decreases as *P*_sig_ increases (Fig.S4(D)). Intuitively, high levels of signaling causes genes in a 1 state to ‘spread out’, effectively leading to a homogenization of cell-types. The sharp decrease in the number of nodes in response to increase in *P*_adj_ indicates that a low level of inter-cellular connectivity is sufficient for signals to percolate throughout the organism (Fig.S4(E), Fig.S5(B)).

We compared the properties of lineage graphs in our data with those expected for Erdos-Renyi random graphs (ER graphs) of similar size. In ER graphs,there is a fixed probability, *p* of any two nodes in the graph being connected by an edge (Erdös and Rényi, 1959). Therefore, on average, the number of edges in a graph with *n* nodes is proportional to *n*^2^. Increasing the number of nodes to *c* * *n* increases the number of edges to *c*^2^ * *n*^2^. We determined the number of edges in lineage graphs with *n* = [1, 2, 3, 4, 5, 6, 7, 8, 9, 10] nodes and calculated the number of edges expected in graphs with twice the number of nodes if these were ER graphs. We then compared the number of edges in our data with *n* = [2, 4, 6, 8, 10, 12, 14, 16, 18, 20] nodes with what we would expect for ER graphs. Compared to ER graphs, the rate of growth of number of edges in lineage graphs in our data is noticeably slower (Fig.S5(A)).

The number of nodes in lineage graphs decreases sharply with the parameter *P*_adj_ (Fig.S4(E)). We show here that this occurs because even at low values of *P*_adj_, cell-types in organisms are connected enough that the fraction of cell-types receiving all signals produced in the organisms reaches a maximum (Fig.S5(B)).

The effect of the parameter *P*_asym_ on the number of nodes in lineage graphs can be explained in terms of its effect on the number of distinct daughter cell-types produced (Fig.S5(C)). The number of distinct daughter cells produced at different values of *P*_asym_ is related to the average fraction of genes in a 1 state in these daughter cells. Among all possible cell-types with *N* genes, most cell-types tend to have about half their genes in a 1 state, and very few cell-types contain fewer, or more genes in a 1 state (Fig.S5(D:inset)). Therefore, when 0.2 < *P*_asym_ < 0.6, where on average, daughter cells have about half their genes in a 1 state, organisms produce the most number of distinct daughter cells, and at *P*_asym_ < 0.2 and *P*_asym_ > 0.6, fewer distinct daughter cells are produced (Fig.S5(D)).

**Figure S4:**
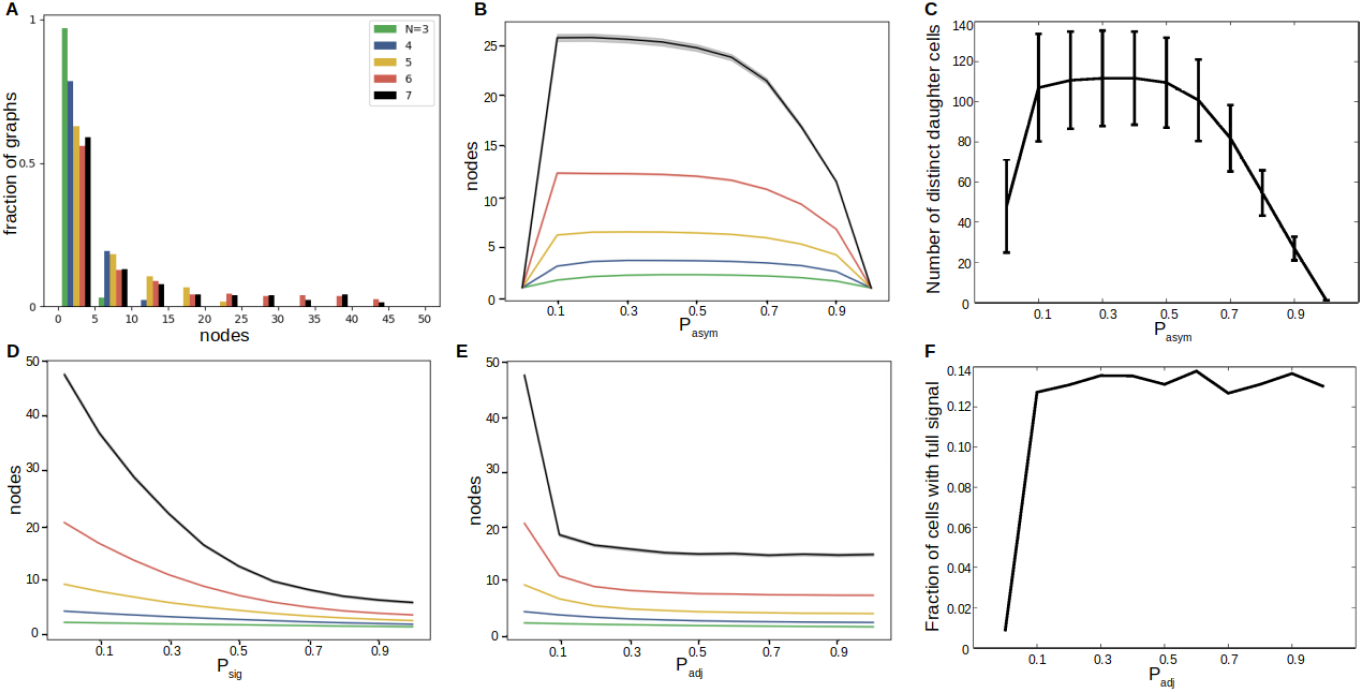
Effect of parameters on lineage graph sizes. **(A)** Histogram of number of nodes in lineage graphs obtained with different N. Histogram bins are of size 5. **(B)** Number of nodes in lineage graphs obtained at different *N* as a function of *P*_asym_. **(C)** Average number of distinct daughter cells produced in an organism as a function of *P*_asym_. At each value of *P*_asym_, 10,000 ‘organisms’ with *N* = 7 genes, composed of randomly chosen cell-types were used to generate these graphs. Error-bars indicate standard deviation. **(D)** Number of nodes in lineage graphs obtained at different *N* as a function of *P*_sig_. **(E)** Number of nodes in lineage graphs obtained at different *N* as a function of *P*_adj_. **(F)** Effect of *P*_adj_ on signal reception. At each value of *P*_adj_, 1000 random signaling vectors SG for *N* = 7 organisms, generated at *P*_sig_ = 0.5 were used. In each organism, the set of signals received by a randomly chosen cell-type, from all *2^N^* possible cell-types in the system was assessed. The horizontal axis represents *P*_adj_, and the vertical axis represents the fraction of cell-types out of 1000, that received all possible signals. In **(B,D,E)** thick lines represent the mean and shaded regions around the lines represent standard deviation (the shaded regions are hard to see because the standard deviations are low). 4852994 graphs were used to generate plots in **(A,B,D,E)**.

### 1.4 Lineage graph topologies and graph randomization

Lineage graphs can be either cyclic or acyclic. The acyclic graphs can be further classified into (i) chains, (ii) trees (acyclic graphs with branches) and (iii) DAGs (Directed Acyclic Graphs, which contain edges connecting different branches). And cyclic lineage graphs can be further classified into (i) unicellular (single cell-type), (ii) SCC (Strongly Connected Component – all paths are cyclic), and (iii) cyclic (contains both cyclic and acyclic paths), (iv) chains (acyclic graphs with no branches) (Fig.S6(A)).

Although all topologies are spread widely across parameter space, different topologies are enriched in different regions of parameter space (Fig.S6(B,C)). No parameter region is monopolized by a single topology, except at extreme values of *P*_asym_, where, as discussed earlier, most graphs are unicellular.

In our data, the most sparsely sampled ingredient is genome regulation. Thus, we wanted to find out the extent to which graph topology distributions (i.e. fractions of all graphs which are unicellular, cyclic/SCC, chains, trees or DAGs) depend on the details of genome regulation matrices we use. To do this, we compare lineage graph topology distributions produced by bootstrap samples to those in the full data. The topology distributions were obtained by pooling all graphs produced in the sample across all parameters. The analysis indicates that graph topology distributions do not depend strongly on the details of the genome, and even bootstrap samples one-fourth the size of the original data produce similar topology distributions (Fig. S7).

**Figure S5.**
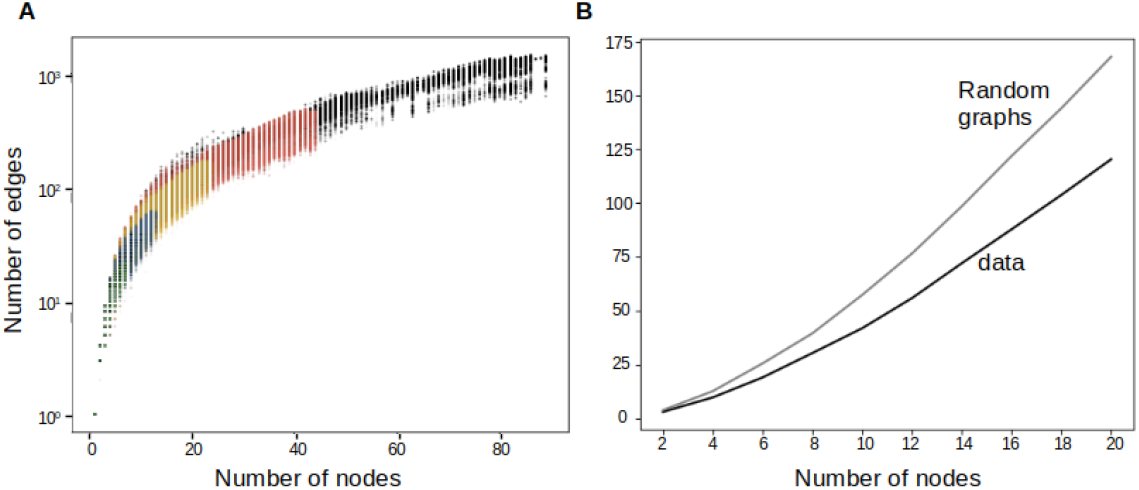
Lineage graphs versus Erdos-Renyi random graphs. **(A)** scatter plot of number of edges and number of nodes in lineage graphs. Transparency has been added to points to make density of points more apparent. 4852994 graphs were used to generate this plot.**(B)** A comparison of growth rate of the number of edges with number of nodes in lineage graphs of our data, versus that expected of Erdos-Renyi random graphs. 4532110 graphs were used to generate this plot.

To a large extent, lineage graph topologies can be characterized by their in-degree and out-degree distributions. For instance, in chains, in-degrees and out-degrees are at most 1, whereas in SCCs, in-degrees and out-degrees are at least 1 (Fig.S8(A)). Therefore, for the most part, we can explain the model’s propensity to generate certain topologies, in terms of its propensity to generate certain in-degree and out-degree distributions.

We randomized lineage graphs generated with our model while keeping node in-degrees and out-degrees unchanged. Topology distribution largely remains unchanged upon randomization (Fig.S8(C,D)), and not many graphs change their topology upon randomization (Fig.S8(B)). Although, we find that the proportion of acyclic graphs decreases slightly, from 24% in model generated graphs, to 19% in randomized graphs.

### 1.5 Characteristics of lineage graphs with different topologies: graph size

Different graph topologies are different in their graph size distributions. While SCC and cyclic graphs span a large range of graph sizes (Fig.S9(A,B)), trees and chains tend to be notably small (Fig.S9(C,E)). DAG type graphs can have moderately large number of nodes (Fig.S9(D)).

### 1.6 Characteristics of tree-type lineage graphs

Tree-type graphs can be further characterized as divergent or convergent trees. We call graph nodes with in-degrees > 1 convergent, and nodes with out-degrees > 1 divergent. Note that by this definition, the same node is allowed to be both convergent and divergent. For some tree-like graph with *n* nodes and *n_e_* edges, let *in_i_* and *out_i_* be the in-degree and out-degree of the *i^th^* node respectively. We define for this graph a number *g_c_* as the sum of in-degrees of all convergent nodes, and a number *g_d_* as the sum of out-degrees of all divergent nodes, i.e.;

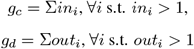

**Figure S6:**
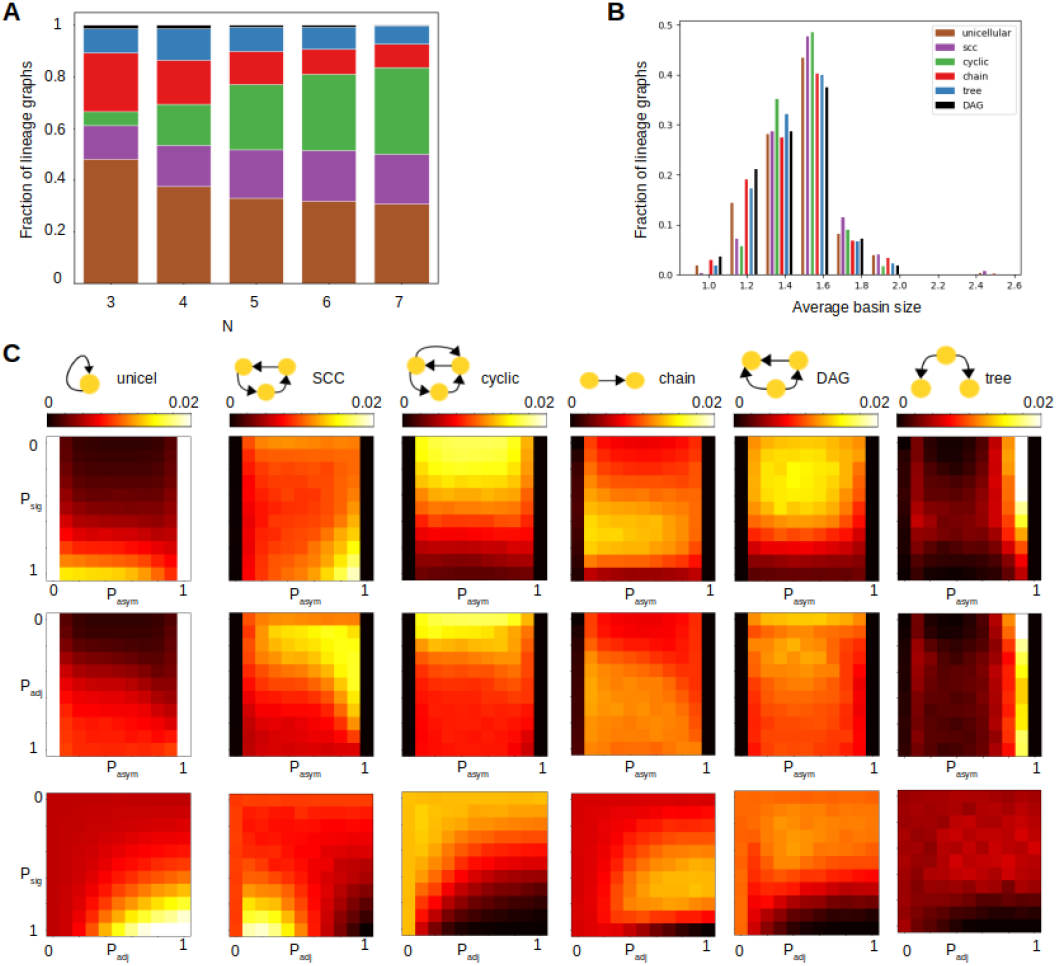
Lineage graph topologies. 4852994 graphs were used to generate these plots. **(A)** Stacked histogram for topologies of lineage graphs obtained with different N. Different topologies are represented with different colours: unicellular:brown, SCC:purple, cyclic:green, chain:red, DAG:blue, tree:black. Heights of colored blocks represent the proportions of corresponding topologies. **(B)** histogram of average basin sizes in the gene regulatory networks of homeostatic organisms with lineage graphs of different topologies. **(C)** 2-D histograms indicating distribution of topologies across parameter space. The first row of histograms show distributions along *P*_asym_ and *P*_sig_, second row along *P*_asym_ and *P*_adj_, and the third row along *P*_adj_ and *P*_sig_. Different columns correspond to histograms for different topologies, as indicated at the top of each column. Intensity of colours in histograms in any column indicates the fraction of graphs of a particular topology found in the corresponding parameter region, according to the colorbars given at the top of each column.

**Figure S7:**
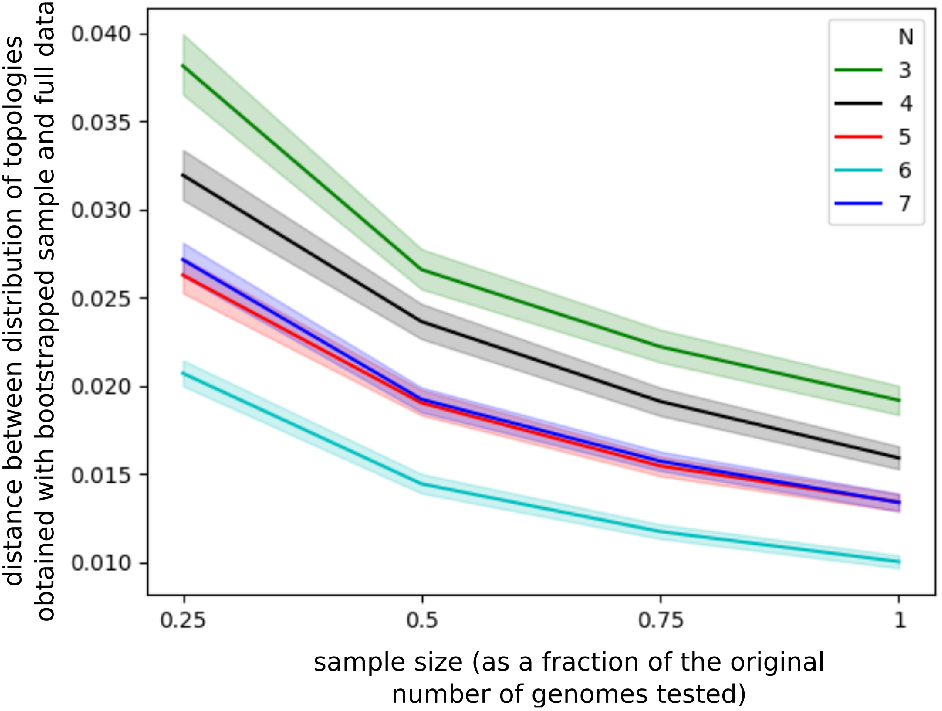
Effect of genome regulation. 4852994 graphs were used to generate this plot. Euclidean distance between normalized topology distributions obtained with bootstrap samples and those obtained with the full data for each N. We tested samples with 25%, 50%, 75% and 100% of all *GR* matrices used in the data for a given N. 1000 samples were generated for each sample size. Lines indicate mean distance and shaded region indicate 95% confidence intervals.

We define the degree of divergence of this graph as (*g_d_ — g_c_*)/*n_e_*. For a perfectly divergent tree, such as the tree to the left in Fig.S10(A), the degree of divergence is 1. And for a perfectly convergent tree (e.g. the tree to the right in Fig.S10(A)), degree of convergence is −1. We find that most tree-like graphs in our data tend to be more convergent than divergent (Fig.S10(B)). Lineage graphs that are divergent lead to an increase in cell-type diversity starting from a few initial cell-types. Lineage graphs of real organisms are believed to be divergent trees. Larger trees tend to be more divergent (Fig.S10(C)). Degree of divergence decreases as *P*_asym_ increases, it is relatively insensitive to *P*_sig_ and *P*_adj_.

### 1.7 Characteristics of DAG-type lineage graphs

DAG-type graphs differ from tree-like graphs in having edges that link different branches. If the edges in the DAG are rendered undirected, these edges are parts of cycles, or loops (Fig.S11(A)). The number of such edges in DAGs can be determined by subtracting the number of edges in the spanning tree of the graph from the total number of edges. For a graph with *n* nodes, the spanning tree has *n* — 1 edges. For a given DAG-type graph, we call the fraction of its edges that forms loops, its link-fraction. Link-fractions of DAG-type lineage graphs indicate the level of trans-differentiation. DAG-type graphs in our data have high link-fractions (Fig.S11(B)), and link-fraction increases with graph size (Fig.S11(C)).

### 1.8 Distribution of regenerative capacities

. In order to infer whether a lineage graph is regenerative, we only look at whether its regenerative capacity is greater than 1, or not. In Fig.S13(A), we show the spread of regenerative capacities for different topologies. For most topologies, median regenerative capacity is greater than 1. The actual value of regenerative capacity is less meaningful, except in the case of tree-type graphs, where most trees have a regenerative capacity of 0. This implies that most trees contain no pluripotent cells. We also find that while median regenerative capacity decreases with *N*, the range of regenerative capacities increases with *N* (Fig.S13(B)).

**Figure S8:**
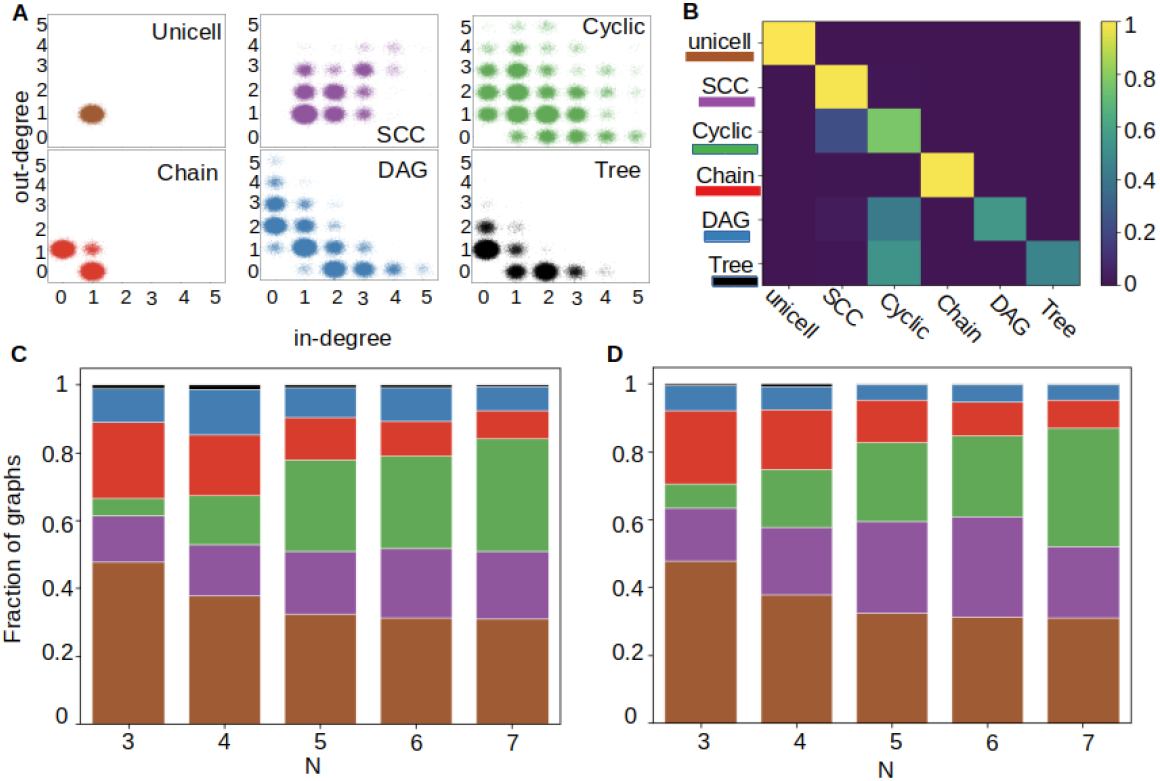
Distribution of topologies of randomized graphs. **(A)** Scatter plots for in-degrees and out-degrees of graph nodes in different topologies. Different topologies are represented by different colours: unicellular: brown, SCC: purple, cyclic: green, chain: red, DAG: blue and tree: black. Noise has been added to points in the plots to make the density of points at each position more apparent. 4852994 graphs were used to generate this figure. **(B)** 2-D histogram representing conversions of graph topology due to randomization. Rows indicate the topologies of original graph and columns indicate the topologies of randomized versions. Intensity of colours in the histogram indicates the fraction of conversions of each type, according to the colorbar given alongside. **(C,D)** Stacked histograms of graph topologies. Heights of coloured blocks indicate the proportion of graphs of the corresponding topology. **(C)** lineage graphs generated by the model, **(D)** randomized lineage graphs. 2373473 graphs were used here.

### 1.9 Intrinsically independent cell-types are enriched in lineage graphs

We wondered whether the large number of independent cell-types in lineage graphs in our data could be attributed to an insensitivity of these cell-types to signals from other cell-types. Alternatively, these cell-types could be independent despite being responsive to signals from other cell-types. We find that the former case tends to be true. We call a cell-type *intrinsically independent* if the full set of signals that can potentially be received by each of its daughter cells is already satisfied by signaling among these daughter cells themselves. In other words, no further external signals can influence the fates of the daughter cells of intrinsically independent cell-types. We calculated the fraction of intrinsically independent cell types across all 2^*N*^ possible cell-types across all systems in our data. We find that cell-types that are part of lineage graphs are much more likely to be intrinsically independent irrespective of parameter region (Fig.S14). Thus cell-types in lineage graphs are predisposed to be independent. But, not all independent cell-types in lineage graphs are intrinsically independent (overall, about 20% the independent cells across all lineage graphs are not intrinsically independent), and not all independent cell-types are pluripotent (Fig.S14(A)).

In lineage graphs, most pluripotent cells are independent of cellular context (Fig.S15(B)). Interestingly, while the proportion of independent cell-types pooled from all graphs is similar (≥ 75%) across all topologies, different topologies have very different proportions of pluripotent cells (Fig.S15(A), fourth panel). Notably, in SCC-type lineage graphs, where all differentiation paths are cyclic, 99.8% of all independent cells are pluripotent. Whereas in lineage graphs that contain acyclic differentiation paths, the proportion of pluripotent independent cells is lower; particularly in tree-type lineage graphs, where only 2.4% of the independent cells are pluripotent. More generally, this indicates that not only the number of independent cell-types, but also their connectivity in the lineage graph is an important factor contributing to an organism’s regenerative capacity.

**Figure S9:**
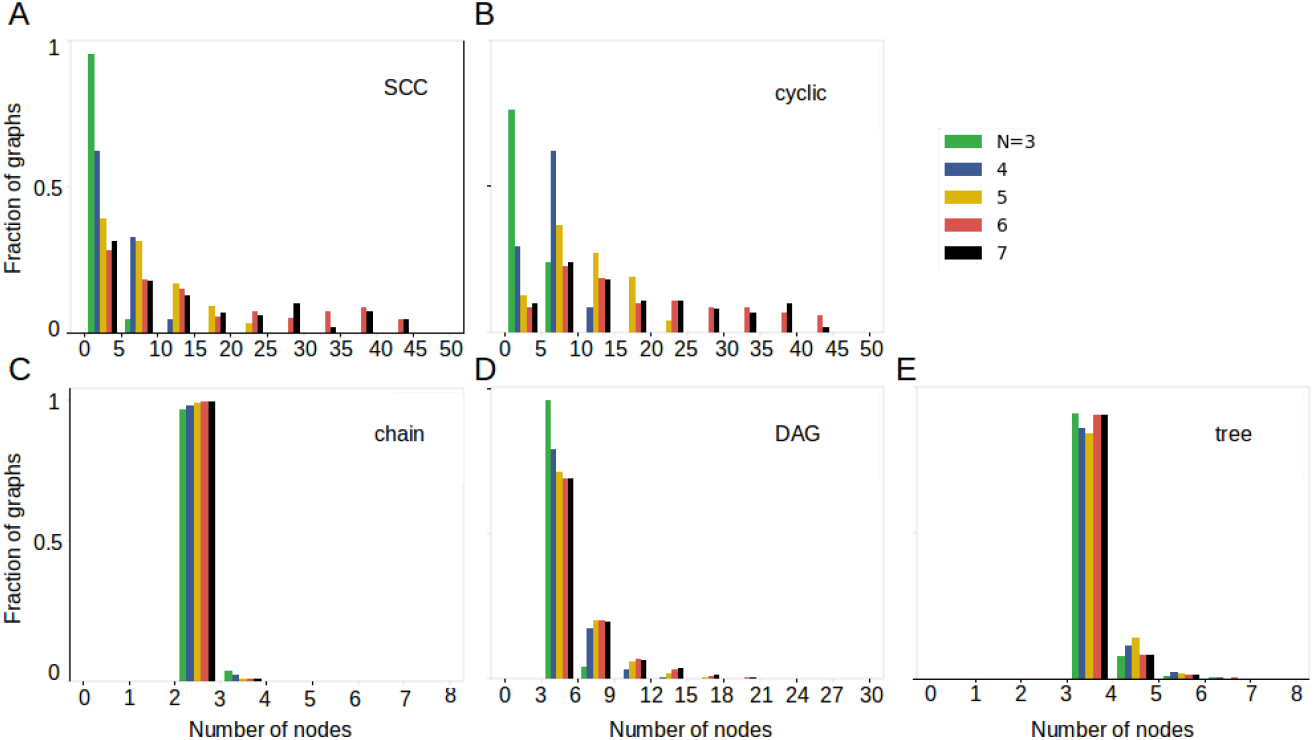
Graph size distributions for different topologies. The horizontal axis indicates number of nodes in lineage graphs and the vertical axis indicates the normalized frequency of graphs. Histogram bin sizes are as follows: **(A,B)** 5, **(C)** 1, **(D)** 3, **(E)** 1. Number of graphs used in these figures are: **(A)** 850101, **(B)** 1044178, **(C)** 695972, **(D)** 474582, **(E)** 46554.

### 1.10 Rules with higher levels of signaling generate lineage graphs with non-root pluripotent cells

We looked at the distribution of parameters *P*_asym_, *P*_sig_, *P*_adj_ of developmental rules encoding lineage graphs with only pluripotent root nodes, and those with non-root pluripotent nodes. We see no significant difference in the distribution of *P*_asym_, that generate the two kinds of lineage graphs (Fig.S16(A,D)). But, consistently with our arguments, we find that lineage graphs with non-root pluripotent cells are more likely to be produced by rules with increased signaling (Fig.S16(B,C,E,F)).

### 1.11 Cell death may increase the proportion of acyclic lineage graphs

Although cells do not undergo programmed death in the current version of the model, here we demonstrate one possible way of implementing programmed cell death, and its implications for lineage graph topologies. We consider ‘cell-death’ as an additional stable cell-state, and a randomly chosen set of cell-types are assigned to its basin. We implement cell-death concomitantly with the gene regulation step in the current version of the model. Cell-types that map to ‘cell-death’ are removed from the organism and not carried forward to the updated state of the organism. We find that allowing some cell-types to die, even though we start from the same initial condition, and use identical rules for signaling and cell-division, can result in completely different homeostatic organisms. Importantly, the nodes of these lineage maps are no longer bound to have out-edges (Fig. S17). Such nodes should increase the propensity of the model to produce acyclic graphs.

**Figure S10:**
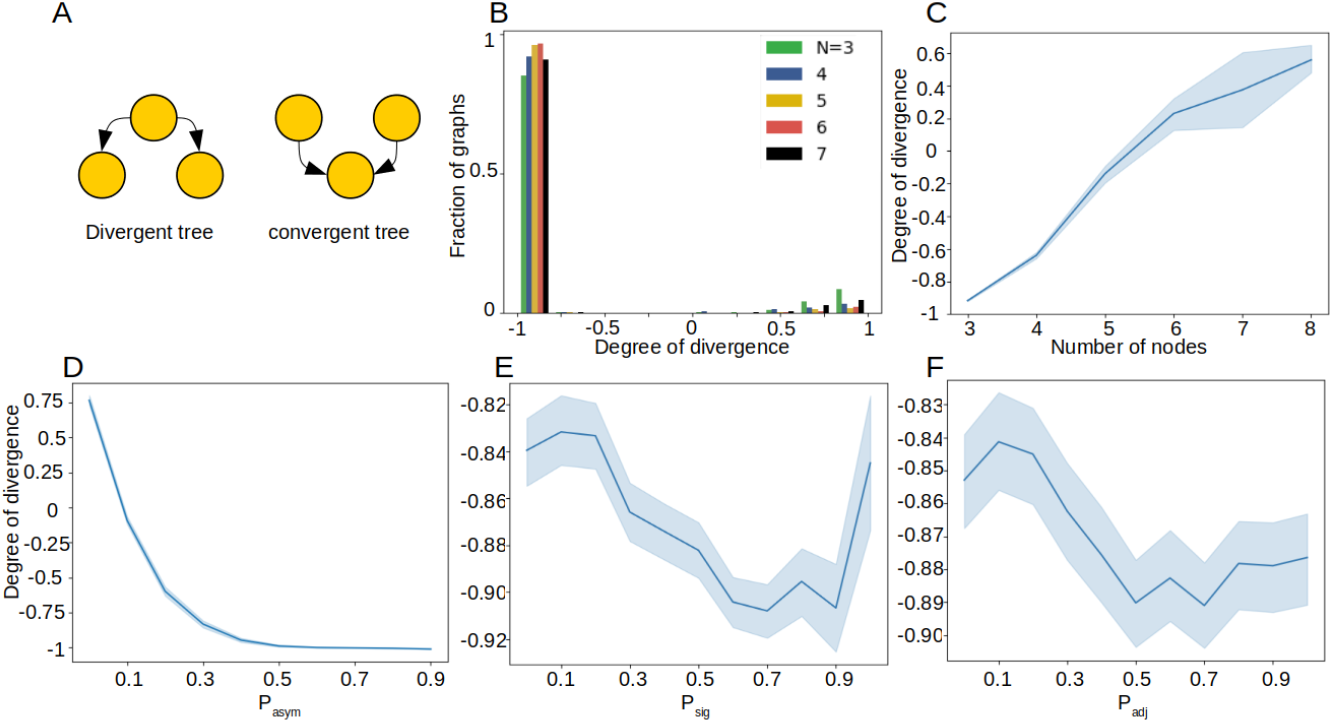
Properties of tree-type graphs. 46554 graphs were used to generate these plots. **(A)** Schematics of a divergent tree like lineage graph and a convergent tree like lineage graph. Yellow circles represent cell-types and edges represent lineage relationships. **(B)** Histogram of degrees of divergence for tree-like graphs in our data with different N. **(C,D,E,F)** Average degree of divergence in our data as a function of **(C)** number of nodes in lineage graphs, **(D)** *P*_asym_, **(E)** *P*_sig_, **(F)** *P*_adj_. Shaded regions indicate standard deviation.

### 1.12 ‘Acyclized’ versions of cyclic graphs

We derived ‘acyclized’ versions of cyclic graphs in our data by merging all nodes belonging to any SCC (strongly connected component) into a single node. In the acyclized graph, there is an edge from some node A to some node B, if in the original graph there is at least one node in SCC A that gives an edge to at least one node in SCC B. Among these acyclized graphs, 45% are unicellular, i.e. the original graphs had a single SCC. DAGs are the next most abundant acyclized graphs, and trees are the least abundant (Fig. S18A). In order to measure *how cyclic* any graph is, we use two metrics: *acyclic edge fraction*, and *acyclic node fraction* which are the ratios of (number of edges/nodes in the acyclized graph) / (number of edges/nodes in the original graph). For the large fraction of acyclized graphs that are unicellular, evidently, both measures are 0. For non-unicellular acyclized graphs, the acyclic edge fractions tend to be very small (on average 0.29), while acyclic node fraction is on average 0.5 (Fig. S18B). In Fig. S18C, we calculate the fraction of trees across all graphs in the data using relaxed definitions for tree-type graphs. We find that even after substantially relaxing constraints and counting all cyclic graphs with acyclic edge fraction ≥ 0.5 as acyclic graphs, and counting all DAGs with *n_l_*/*n_b_* ≤ 0.5 as trees, the fraction of trees across all graphs remains very low. We also check whether relaxing the definition of trees effects our results on the regenerative capacities of different topologies (Fig.S19). Although the median fraction of regenerative tree-type graphs increases from 0.1232 to 0.1911, qualitatively, this fraction is still the lowest among all topologies considered.

**Figure S11:**
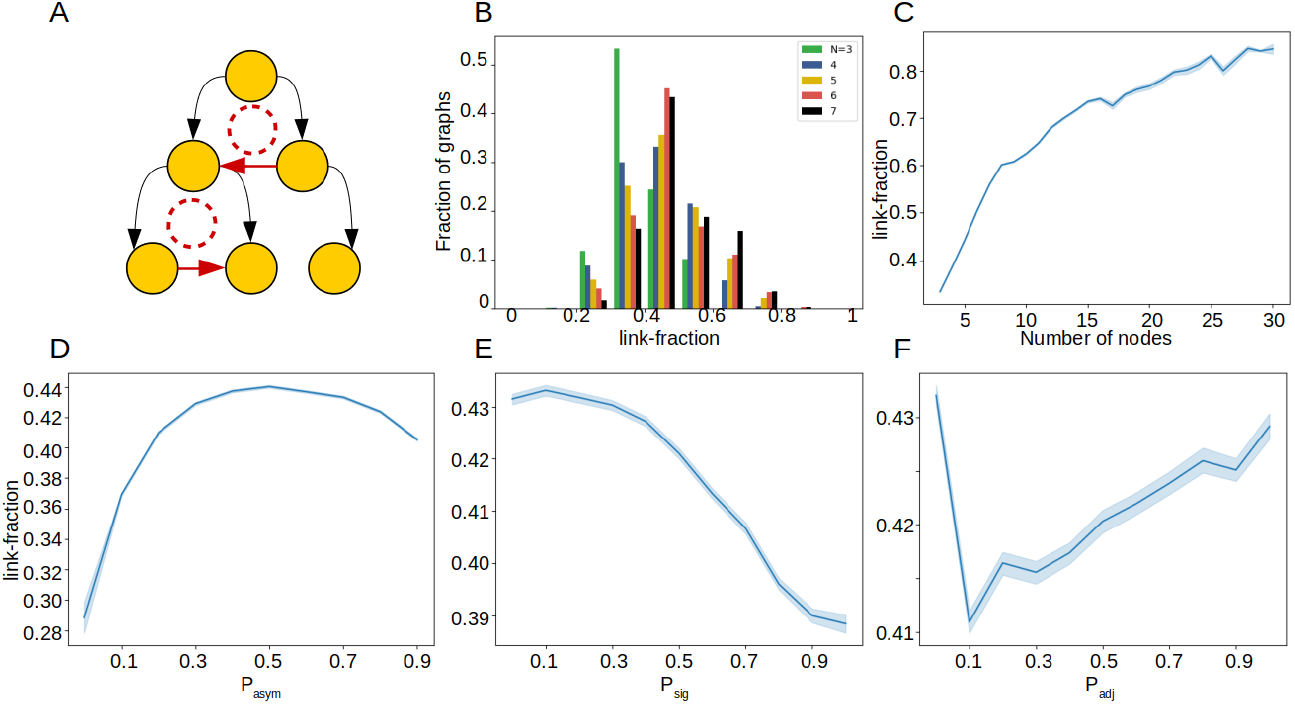
Properties of DAG-type graphs. 474582 graphs were used to generate these plots. **(A)** Schematic of DAGs. Yellow circles represent cell-types, and edges represent lineage relationships. Red edges forms loops in the DAG. **(B)** Histogram of link-fraction of DAGs in our data. Loop-fraction is defined as the fraction of edges in a DAG that form loops. Histogram bins are of size 0.1. **(C,D,E,F)** Average loop-fraction of DAG type graphs in our data as a function of **(C)** number of nodes in lineage graphs, **(D)** *P*_asym_, **(E)** *P*_sig_, **(F)** *P*_adj_. Shaded regions indicate standard deviation.

**Figure S12:**
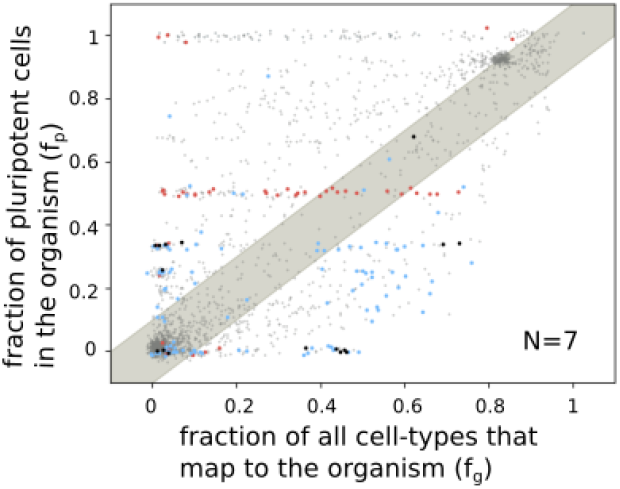
Regenerative capacity and isomorphic graphs. 13177 graphs were used to generate this plot. Here, we plot the same graph as in Fig3(A) in the main paper, but only include points for graphs that are not isomorphic to any other graph with the same values of *f_g_* and *f_p_*. This reduces the density of points at some of the ‘clusters’ which appear in Fig4(A).

**Figure S13:**
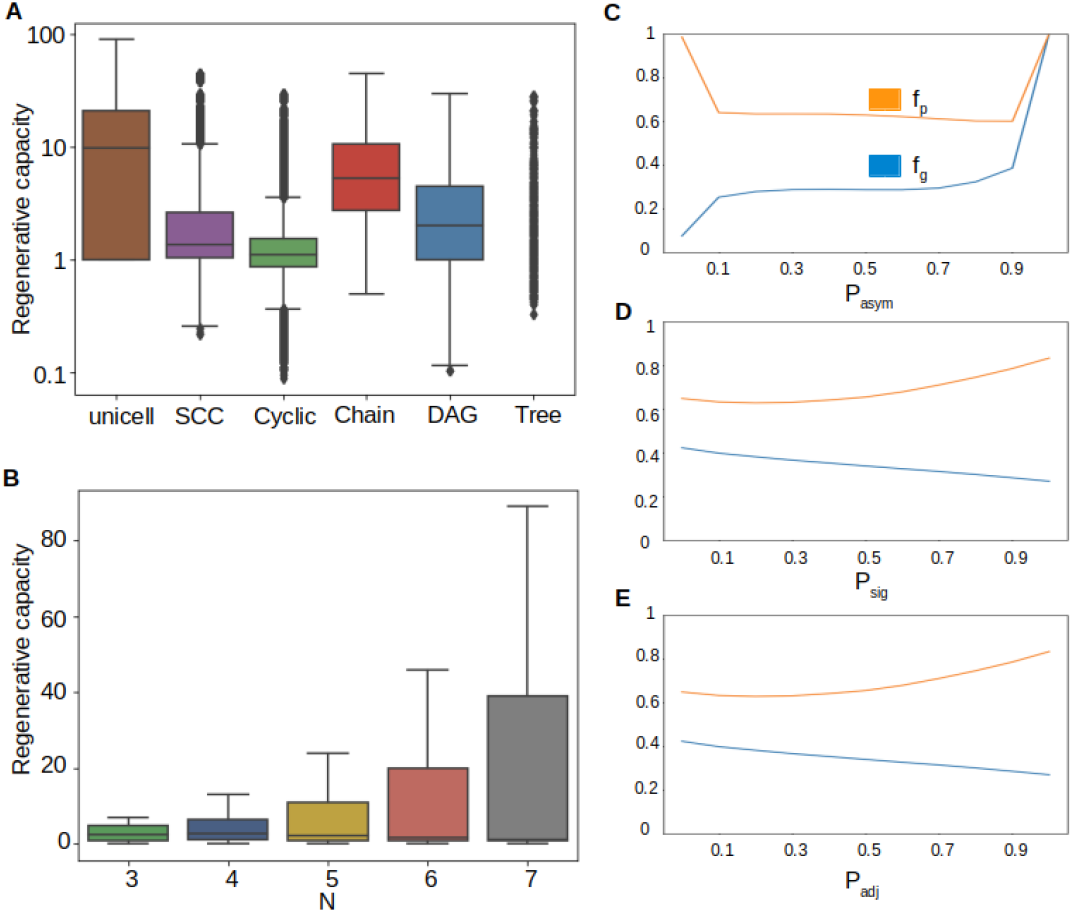
Box plots for regenerative capacity of lineage graphs. 4852994 graphs were used to generate these plots. **(A)** across different topologies, **(B)** across number of genes N. Boxes represent quartiles of the data set. Lines inside the box shows the median, while whiskers show the rest of the distribution. Outliers are shown as diamonds. Most tree-like lineage graphs have a regenerative capacity of 0, therefore the box for these graphs is not visible.**(C,D,E)** Variation of regenerative capacity across model parameters: **(C)** *P*_asym_, **(D)** *P*_sig_, **(E)** *P*_adj_. Fraction of pluripotent cells (*f_p_*) is shown in orange, and the fraction of all cells (present or absent from organisms) that develop into the organism (*f_g_*) is shown in blue. Bold lines represent mean values (shaded regions around the lines represent standard deviations, which are small and hardly noticeable). The average regenerative capacities of graphs at different parameter values can be judged by the difference in the heights between the orange and blue curves.

**Figure S14:**
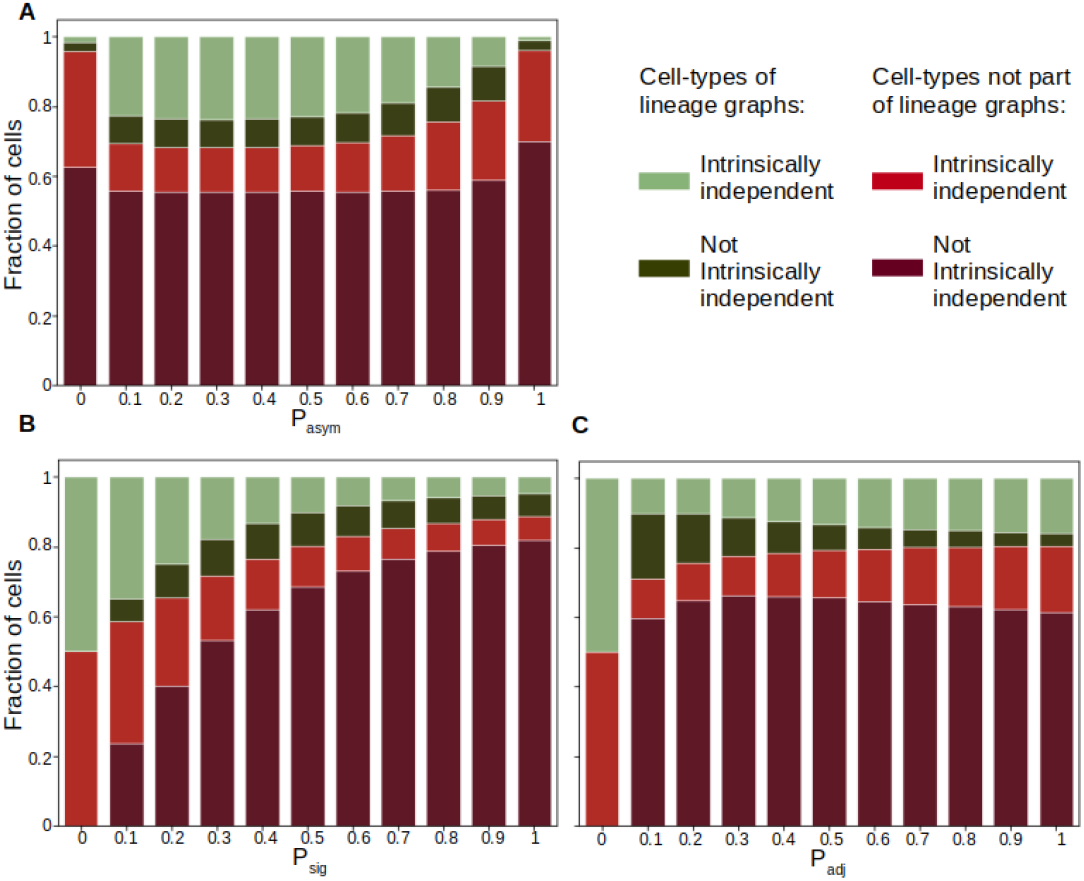
Stacked histograms showing intrinsic independence of cell-types. 4852994 graphs were used to generate these plots. **(A)** as a function of *P*_asym_, **(B)** as a function of *P*_sig_, **(C)** as a function of *P*_adj_. Different cell-type categories are represented with different colours. Cell-types not part of organisms are represented in reds; intrinsically independent: bright red, not intrinsically independent: dark red. Cell-types found in organisms are represented in greens; intrinsically independent: light green, not intrinsically independent: dark green. Heights of colored blocks represent the proportions of corresponding cell-types.

---

**Figure S15:**
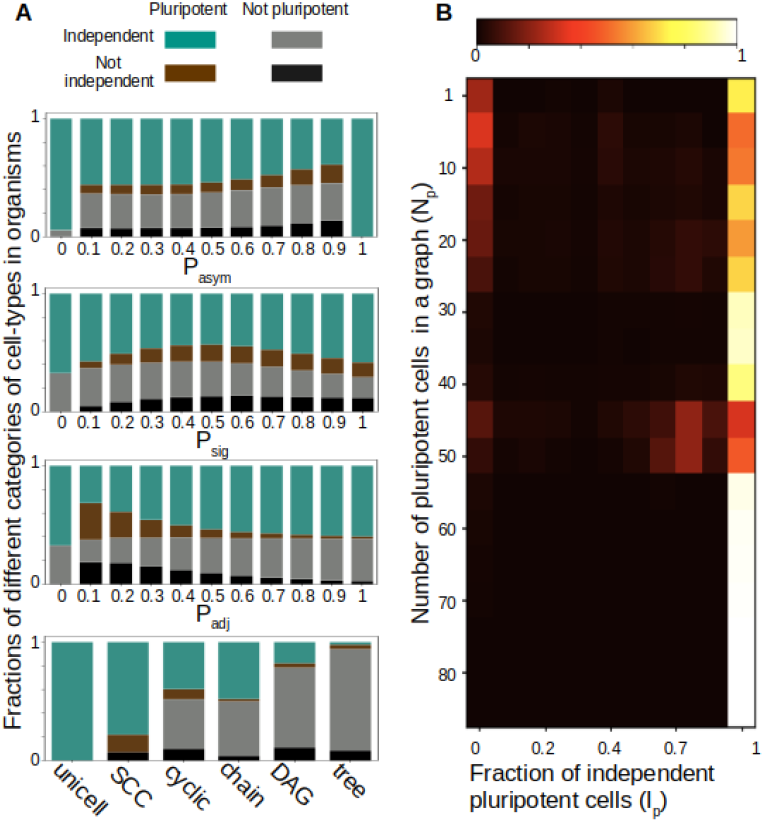
Independent pluripotent cell-types. 4852994 graphs were used to generate these plots. **(A)** Stacked histograms for cell-types of different categories pooled from organisms across different parameter values (top 3), or across lineage graphs with different topologies (see also Fig.S1, Fig.S2). Different cell-type categories are represented with different colours. Non-pluripotent cells are represented in greys; independent: light grey, not independent: black. Pluripotent cells are represented in colours; independent: teal, not independent: brown. Heights of colored blocks represent the proportions of corresponding cell-types.**(B)** 2-D histogram indicating the fraction of independent pluripotent cell-types in homeostatic organisms. Intensity of colours in the histogram indicate the fraction of organisms with *N_p_* pluripotent cell-types, *I_p_* of which are independent, according to the colorbar given on top.

**Figure S16:**
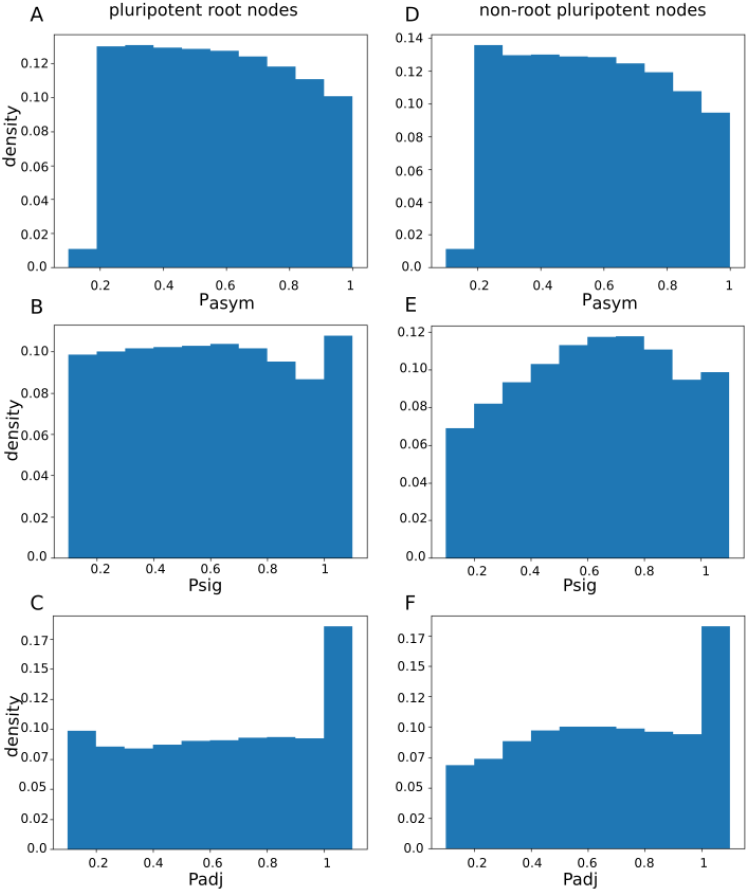
Comparison of parameters that generate regenerative acyclic lineage graphs with pluripotent root nodes versus those with non-root node pluripotent cells. **(A,B,C)** Parameter distributions for graphs with pluripotent root nodes. 663849 graphs were used to generate these plots. **(D,E,F)** Parameter distributions for graphs with non-root pluripotent nodes. 463054 graphs were used to generate these plots.

**Figure S17:**
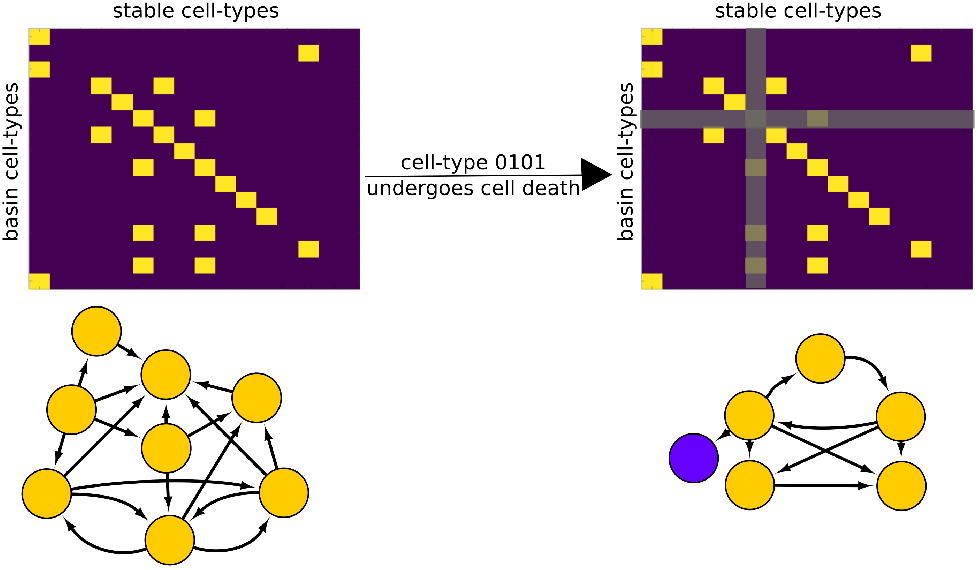
Effect of including cell-death in the model. The matrix on the left represents the *GR* matrix for generating the ‘cyclic’ graph shown in Fig2(B) in the main paper, and the matrix on the right represents the modified *GR*, where cell-type 0101 undergoes cell-death. Below the matrices are the lineage graphs generated using the corresponding *GR* matrices, while the initial condition, and the rest of the rules matrices are kept the same as those used to generate the original ‘cyclic’ lineage map. In the lineage graph on the right, the purple node has no out-edges (not even a self-edge).

**Figure S18:**
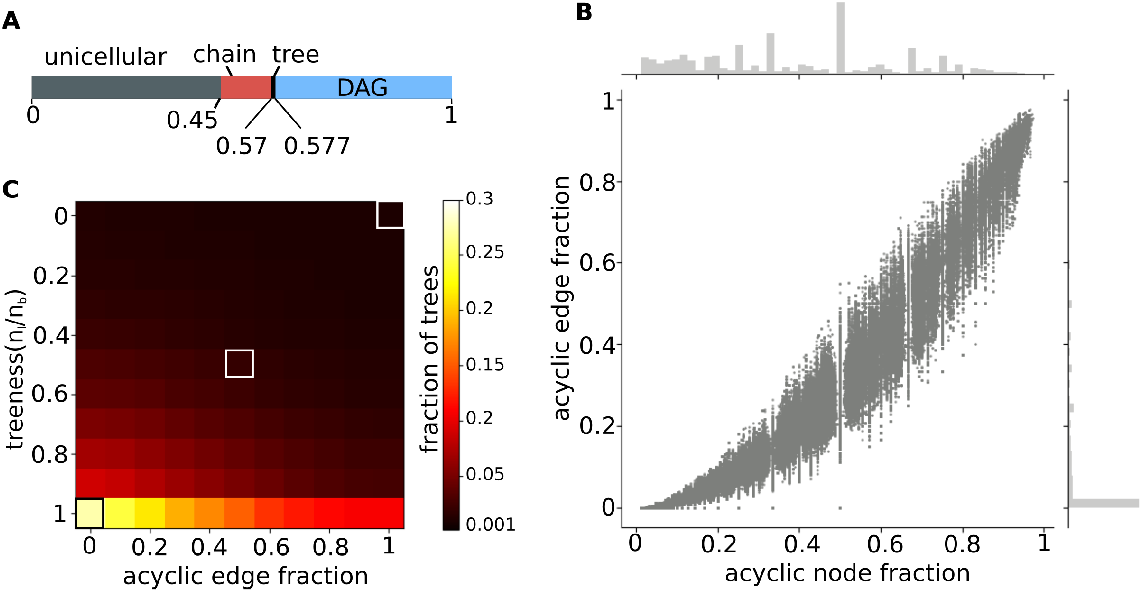
Properties of ‘acyclized’ cyclic graphs. 1894279 graphs were used to generate **(A,B)**. **(A)** Topology distribution of ‘acyclized’ lineage graphs. **(B)** Scatter plot of *acyclic node fraction* versus *acyclic edge fraction* for non-unicellular acyclized lineage graphs. The histograms opposite the axes represent the marginal distributions for the corresponding axes. **(C)** Heat-map for the fraction of trees across all graphs in the data when the definitions of acyclic graphs and tree-type graph are relaxed. The *acyclic edge fraction* threshold for considering a cyclic graph as acyclic relaxes from right to left on the x-axis. The *n_l_/n_b_* threshold for considering a branched acyclic graph as a tree relaxes from top to bottom. The white-edged square on the top-right represents the case where strict definitions for acyclic and tree-type graphs are used (fraction of trees is 0.01). In the white edged square in the middle (edge-fraction threshold = 0.5, *n_l_*/*n_b_* threshold = 0.5), the fraction of tree-type graphs is 0.02. The black-edged square on the bottom-left represents the most relaxed case where all cyclic graphs are considered acyclic and all branched acyclic graphs are considered trees. Here, the fraction of tree-type graphs 0.3. 4852994 graphs were used to generate this plot.

**Figure S19:**
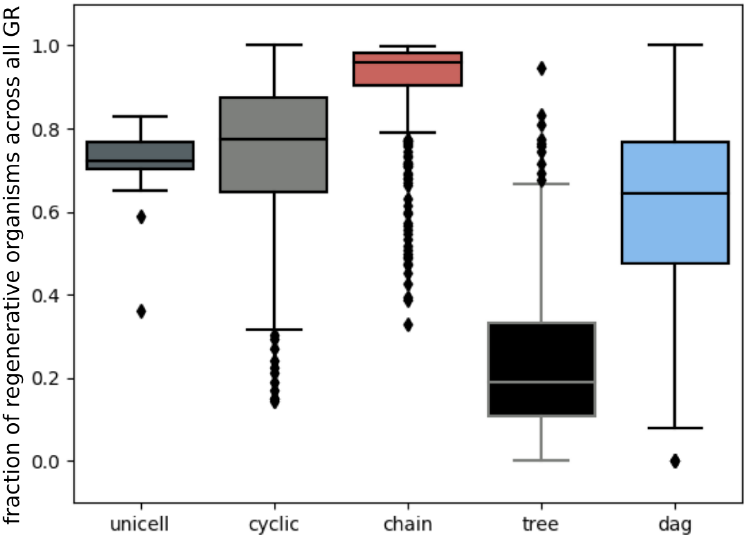
Box plots for regenerative capacities using relaxed definitions for acyclic graphs and trees. Here, we treated acyclized graphs with edge-fractions≥ 0.5 as acyclic graphs. In addition, we treated acyclic graphs with *n_l_*/*n_b_* ≤ 0.5 as trees. For each *GR* used in our data, for a given graph topology, we looked at the fraction of graphs with regenerative capacity > 1 (as described in the main paper). Boxes represent quartiles of the data set. Lines inside the box show the median, while whiskers show the rest of the distribution. Outliers are shown as diamonds. 4852994 graphs were used to generate this plot.

## Notes

### Competing Interest Statement

The authors have declared no competing interest.

### Summary of Updates

Additional tests of the model, in particular, characterization of topologies. Further discussion of possible empirical tests.

